# Small non-coding vault RNA1-2 modulates expression of cell membrane proteins through nascent RNA silencing

**DOI:** 10.1101/2022.08.12.503724

**Authors:** Adele Alagia, Jana Terenova, Ruth F. Ketley, Arianna Di Fazio, Irina Chelysheva, Monika Gullerova

## Abstract

Gene expression can be regulated by transcriptional or post-transcriptional gene silencing. Recently, we described nuclear nascent RNA silencing (NRS) that is mediated by Dicer dependent tRNA-derived small RNA molecules. In addition to tRNA, RNA polymerase III also transcribes Vault RNA, a component of the ribonucleoprotein complex Vault.

Here, we show that Dicer dependent small vault RNA1-2 (svtRNA1-2) associate with Argonaute 2 (Ago2). Whilst endogenous vtRNA1-2 is present mostly in cytoplasm, svtRNA1-2 localises predominantly in nucleus. Furthermore, in Ago2 and Dicer knockdown cells, a subset of genes which are upregulated at the nascent level were predicted to be targeted by svtRNA1-2 in the intronic region. Genomic deletion of vtRNA1-2 results in impaired cellular proliferation and the upregulation of genes associated with cell membrane physiology and cell adhesion. Silencing activity of svtRNA1-2 molecules is dependent on seed-plus-complementary-paired hybridisation features and the presence of a 5-nucleotide loop protrusion on target RNAs.

Our data reveal a role for Dicer dependent svtRNA1-2, possessing unique molecular features, in modulation of expression of membrane associated proteins at the nascent RNA level.

## Introduction

Regulation of gene expression is fundamental for the control of cellular processes such as stemness, differentiation, development, and adaptation to environmental cues [1]. Control of RNA expression at the transcriptional and post-transcriptional level can be achieved by several mechanisms [2]. For example, changes in chromatin accessibility through transcriptional gene silencing (TGS), and mRNA translation inhibition or degradation by post-transcriptional gene silencing (PTGS), can efficiently coordinate the silencing of the cellular transcriptome [3]. RNA interference-related gene silencing mechanisms facilitate specific RNA downregulation with the “guidance” of various classes of small non-coding RNA (sncRNA) molecules such as microRNA (miRNA) [4], small interfering RNA (siRNA), and Piwi-interacting RNA (piRNA) [5]. Furthermore, tRNA-derived small RNA (tsRNA) [6], rRNA-derived fragments (rRFs) [7], snoRNA-derived RNAs, and YRNA-derived fragments have been recently identified as small non-coding regulatory RNA molecules [8].

We have previously identified a class of tsRNA molecules derived from tRNAs, produced by Dicer cleavage activity and loaded onto Ago2, which target RNA intronic sequences resulting in nascent RNA silencing (NRS) [9]. The mechanism of NRS differs from TGS and PTGS as the transcription of the target gene remains unaffected and the nascent RNA is degraded within the nucleus. Although we described the fundamental mechanism of NRS, our understanding of the molecular and functional details remains limited.

In the human genome, vault RNAs are located on chromosome 5q31 within two different loci: the VAULT-1 locus which consists of three genes (VTRNA1-1, VTRNA1-2 and VTRNA1-3), and the VAULT-2 locus which encodes for VAULT2-1 [10]. Both loci are under the control of a RNA Polymerase III type 2 promoter which contains box A and box B motifs [11], typically found within tRNA genes. Vault RNA molecules are 80 to 140 nucleotides long [12, 13] and classified as sncRNAs. VtRNA was initially identified as a component of the large ribonucleoprotein particle known as Vault, which is involved in several cellular processes, such as nuclear transport [14, 15], immune response [16], and drug resistance [17–19]. However, it has been shown that the majority of vtRNA molecules (~95%) are not associated with the Vault particles [20], suggesting that vtRNA molecules might have additional roles within the cell, including as ribo-regulators of autophagy [21] or as effectors of gene expression control in a miRNA-like fashion [22]. Indeed, some vtRNA molecules can be processed into smaller fragments called small vault RNA (svtRNA), and their biogenesis can be Drosha-and Dicer-independent (*i.e*. svtRNA1-1), or Drosha-independent and Dicer-dependent (*i.e*. svtRNA2-1) [22]. Furthermore, svtRNAs biogenesis can be regulated by serine/argine-rich splicing factor (SRSF2) and the deposition of 5-methyl-cytosine modification in an Nsun2-dependent manner. SvtRNA1-1 and svtRNA2-1 molecules can also associate with Ago proteins and regulate the expression of genes involved in several cellular pathways such as drug metabolism and encode voltage-gated calcium channels, and act as tumour suppressors [23].

Here, we show that vtRNA1-2 molecules can be actively processed by Dicer into svtRNA1-2 fragments, associate with Ago2 and localise predominantly in nucleus. Using target prediction tool, MiRanda, we identified 227 unique svtRNA1-2 targets with dominant intronic complementarity among genes upregulated in Ago2 and Dicer KD cells. We validated svtRNA1-2-mediated NRS by assessing the nascent and steady state levels of four predicted target genes. In addition, analysis of chromatin-associated RNA-seq (ChrRNA-seq) in HEK293T vtRNA1-2 knockout cells confirmed the involvement of svtRNA1-2 in NRS of a subset of genes, associated with plasma membrane physiology, cell signalling, and cell adhesion, which phenotypically results in defects in cellular proliferation.

Furthermore, we show that svtRNA1-2-mediated silencing relies on both seed-plus-complementary-paired region hybridisation features and the presence of a 5-nt loop protrusion on target RNA. This study is the first report to describe the mechanism of svtRNA1-2 biogenesis and its regulatory role in nascent RNA silencing.

## Results

### vtRNA1-2 is processed by Dicer into small RNA fragments

In order to characterise fragments derived from vtRNA and to examine the potential role of Dicer in the processing of vtRNA, we performed RNA sequencing of small RNA (sRNA) isolated from HEK293T Wild-Type (WT) and Dicer knockdown (KD) (HEK293 cells with integrated inducible shRNA) (**Fig. S1A**) cells [9]. A principal component analysis (PCA) outlined clear differences between the two conditions and tight clustering of the replicates (**Fig. S1B**). The size distribution of mapped reads showed a distinct Dicer-dependent peak between 22-25nt long, including reads corresponding to mature miRNA (21-23nt) (**Fig. S1C**). The small difference in this peak between WT and Dicer KD is most likely due to the presence of the induced shRNA in Dicer KD sample (used to induce Dicer KD), which might mask the absence of a proportion of mature miRNA. Among the identified sRNAs, the most abundant species were miRNA (making up 47% of the population), followed by tRNA (23%). We also identified the four species of vault RNA (**Fig. 1A** and Supplementary Table 1), with >22nt long fragments. Interestingly, fragments derived from vtRNA1-2 were mostly 22-24nt long, suggesting that they might be Dicer products (**Fig. S1D**). Indeed, moving average (MA) analysis showed that fragments derived from vtRNA1-2 and vtRNA2-1 were significantly down-regulated in Dicer KD (p-adj<0.05), suggesting Dicer is important for their biogenesis (**Fig. 1B** and **Fig. S1E**). Interestingly, we identified more svtRNA1-1 reads upon Dicer KD when compared to WT, whilst the expression of vtRNA1-3 was not significantly altered between these samples (**Fig. 1B**), suggesting that Dicer is only involved in the processing of vtRNA1-2 and 2-1. This data is in agreement with additional analysis of small RNA-seq using a previously published dataset in WT and Dicer KD conditions (GSE55333) [24] (**Fig. S1F** and Supplementary Table 4).

**Figure. 1.**
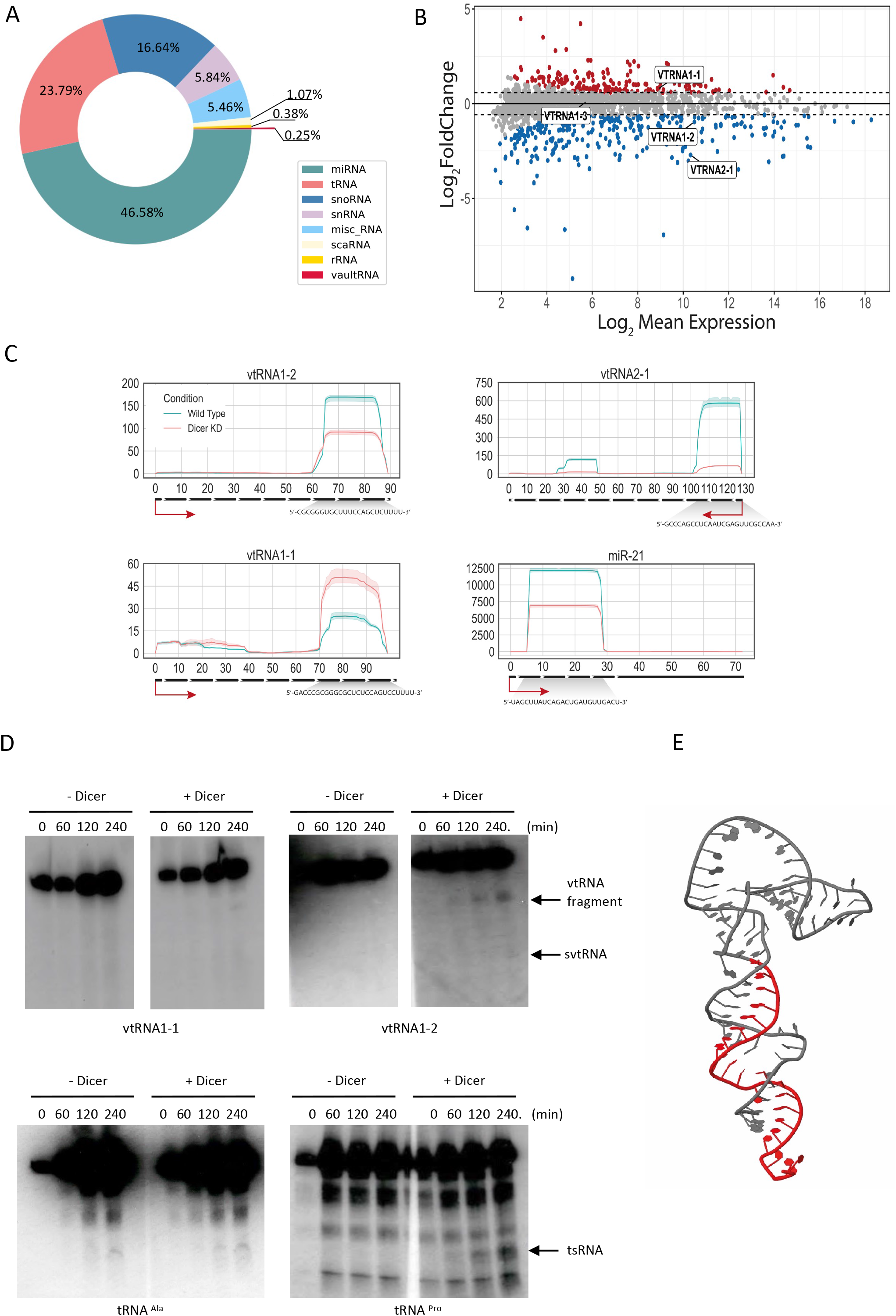
small RNA-seq reveals Dicer processing of vtRNA1-2 into smaller RNA fragments. **A)** Pie chart shows the relative abundance of different RNA species with differentially altered expression in WT and Dicer KD sRNA-seq samples. **B)** MA plot of Dicer KD vs WT. Points in red indicate sRNAs with an adjusted p-value < 0.05 and the points encircled in blue indicate vault RNA. **C)** Per base read coverage of vtRNA. The x-axis shows the length of transcripts, and the y-axis shows the normalised number of mapped reads RPM (Reads Per Million). miR-21 acts as a positive control. **D)** Representative Northern blot images showing signals of vtRNA1-1, vtRNA1-2, tRNA^Pro^ and tRNA^Ala^ at different time points after the addition of purified 6xHis-Dicer. Dicer cleavage products are indicated by arrows. **E)** Predicted tertiary structure of vtRNA1-2. The identified small vault RNA1-2 fragment is coloured in red.

Next, we calculated the per-base read coverage of differentially expressed vtRNA in both WT and Dicer KD conditions. We identified regions along the vtRNA gene sequence with high read coverage, located at the 3’end of vtRNA1-2 and the 5’ end of vtRNA2-1, which were decreased in Dicer KD, indicating that vtRNA1-2 and vtRNA2-1 can be processed into small-vault RNA fragments in a Dicer-dependent manner (**Fig. 1C** and **Fig. S1F**).

To validate our sequencing data experimentally, we have purified recombinant Dicer protein from insect cells (**Fig. S2A and B)** and performed *in vitro* Dicer cleavage assays on vtRNA1-1 and 1-2, using tRNA^Pro^ as positive control, tRNA^Ala^ as negative control [9]. Our data show that Dicer can indeed cleave vtRNA1-2 *in vitro*, although less efficiently that tRNA^Pro^. We did not detect Dicer cleavage activity on tRNA^Ala^ (negative control) and vtRNA1-1 **(Fig. 1D).** It should be noted that the structure of vtRNA1-2 resembles more of a tRNA-like elbow conformation rather than a pre-miRNA molecule with a defined stem loop structure and 2nt 3’overhang, suggesting that vtRNA1-2 is a non-canonical Dicer substrate. Not only is the interaction between the Dicer PAZ domain and the 4nt 3’overhang of vtRNA1-2 not canonical, but the interaction between the Dicer Helicase domain and the vtRNA1-2 stem loop region might also be not be as efficient as in the case of pre-miRNAs. Interestingly, the presence of a hammerhead-like structure in vtRNA1-1 might be a pivotal discriminating factor for Dicer cleavage.

Next, we analysed the reactivity scores from [25] for each nucleotide on the vtRNA1-2 sequence and identified a cluster of highly reactive nucleotides between nucleotides 20 and 50, suggesting that this might be the region of vtRNA1-2 interaction with processing factors (**Fig. S2C**). The newly identified svtRNA1-2 fragment was visualised on the vtRNA predicted secondary and tertiary structure (**Fig. 1E** and **Fig. S2C**). Interestingly, detected reactive nucleotides on vtRNA1-1 were separated into two smaller clusters (around nucleotide 40 and nucleotide 60), which might suggest a difference in processing of vtRNA1-1 and vtRNA1-2.

All four vtRNA sequences are highly similar in the 5’ and 3’ regions but vary in the central domain (between nucleotides 30-60), which could be relevant to the differences in reactive nucleotide distribution (**Fig. S3A**). Interestingly, when we analysed vault RNA1-2 sequence conservation, we confirmed that the 5’ and 3’ regions of vtRNA are highly conserved across vertebrates whilst the middle part is not (**Fig. S3B**). In mice, vtRNA with unknown function is encoded by only one gene, VAULTRC 5, which is longer (143 nt) than any of the human vtRNA. Due to high conservation score, we hypothesised that the processing of vtRNA into svtRNA might be a general evolutionary conserved mechanism. To verify this hypothesis, we analysed a collection of small RNA-seq data from mice (spleen, liver, lungs, muscle, brain, pancreas) (GSE119661). Interestingly, we identified reads localising to a specific region at the 3’ end on the template vtRNA, revealing novel small vtRNA fragments (**Fig. S3C**). Furthermore, we noticed the different expression patterns throughout the analysed tissues, which resembles the tissue-specific expression of vtRNA in humans. These results show the presence of vtRNA-derived fragments in mice tissues, supporting the evidence that svtRNA are not degradation products but instead specific, biologically-functional species. Altogether, these data confirm the existence of an evolutionarily conserved novel Dicer-dependent svtRNA1-2.

### svtRNA1-2 associates with Argonaute proteins and is localised predominantly in nucleus

Next, we set out to test whether svtRNA1-2 is functional. We have previously identified that a group of tsRNA can mediate NRS [9]. As vtRNA share some similarities with tRNA, such as length, Nsun2-dependent m5C modification, Dicer processing, and RNAPIII transcription, we assessed the potential role of svtRNA1-2 in gene silencing. First, we investigated whether svtRNA1-2 fragments associate with Argonaute proteins by analysing publicly available datasets of Ago1-4 (GSE21918) PAR-CLIP (photoactivatable ribonucleoside-enhanced crosslinking and immunoprecipitation), as well as Ago2/3 RIP-Seq (GSE55333) (Supplementary Table 4). Interestingly, we detected a significant enrichment of Dicer-dependent svtRNA1-2 in the Ago2 PAR-CLIP. Fragments derived from vtRNA1-1 were present in the Ago2 PAR-CLIP data only at low levels, and svtRNA1-3 and 2-1 were not detected. We used microRNA-21 (miR-21) as a positive control (**Fig. 2A** and **Fig. S4A**). We also observed svtRNA1-2 in the other Argonaute PAR-CLIP data (Ago1, Ago3 and Ago4) as well as in the Dicer PAR-CLIP dataset (Supplementary Table 4 and **Fig. S4A**).

**Figure. 2.**
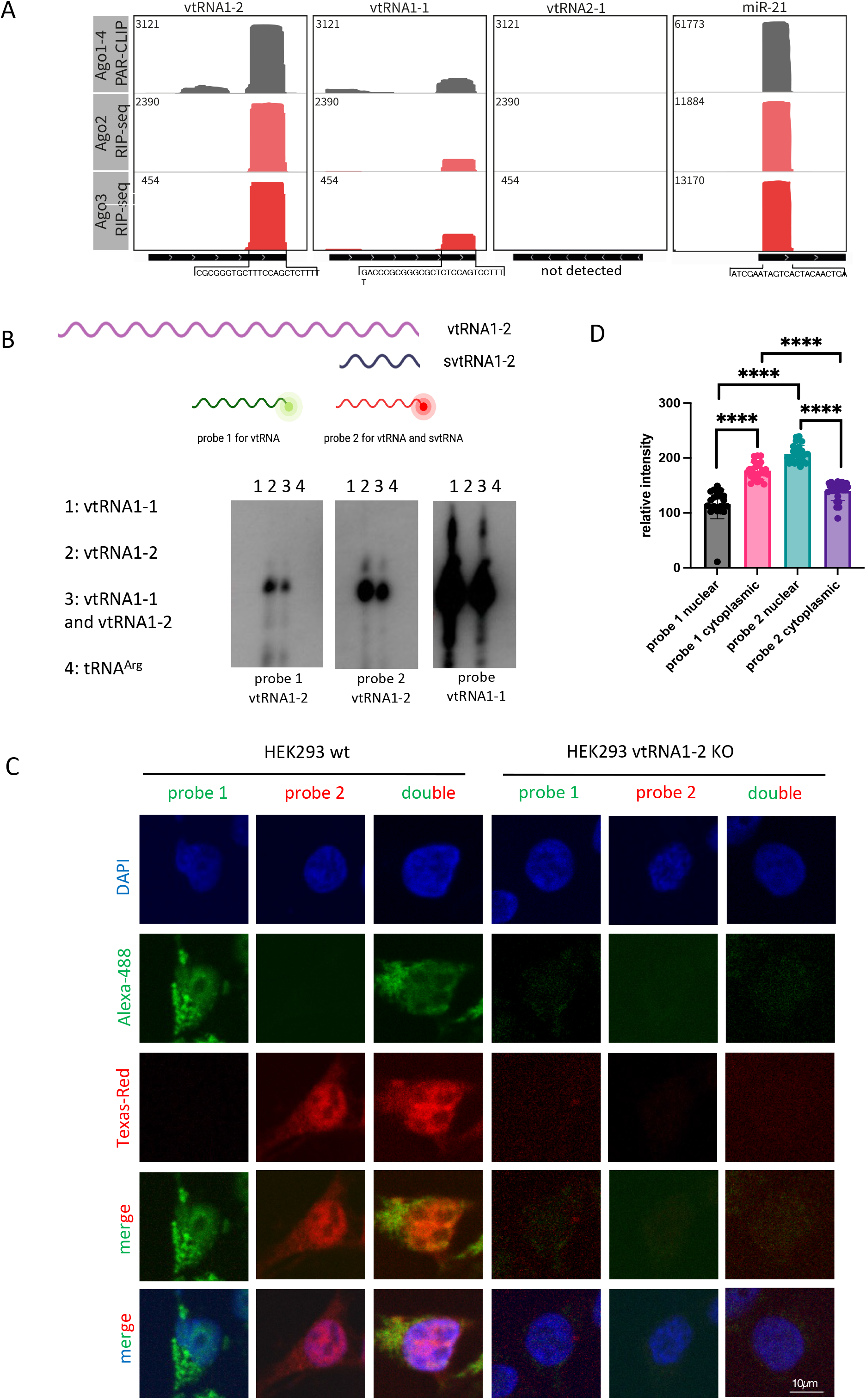
svtRNA1-2 is bound to Argonaute proteins. **A)** Integrative genomics viewer (IGV) tracks of AGO1-4 PAR-CLIP and AGO2/3 RIP-Seq show the association of AGO proteins with svtRNA1-2 (data values are shown in the top left corner). miR-21 was used as a positive control. The bottom black line indicates the length of the transcript and the sequence below represents the interacting fragment. Note: no reads were detected for vtRNA2-1. **B)** Diagram illustrating position of probes 1 and 2 in respect to vtRNA1-2 and svtRNA1-2. Representative Northern blot showing signals for vtRNA1-1 and vtRNA1-2 when incubated with probes 1 and 2 designed for vtRNA1-2. **C)** Representative FISH confocal images showing fluorescent signals for probes 1 and 2 in HEK293 WT and vtRNA1-2 KO cells. DAPI was used to stain nuclei. **D)** Bar chart showing the relative fluorescence intensity levels of probes 1 and 2 in nucleus and cytoplasm. error bar = mean ± SEM; significance was determined using multiple unpaired Student’s *t*-test, \*\*\*\**p*□≤□0.0001.

It has been shown that Ago2, the only member of Argonaute family with well characterized catalytic activity [26], can carry out its function in both the nucleus and the cytoplasm [27–29]. Given that we have shown that tsRNA function in the nucleus [9], we tested whether svtRNA1-2 might be present in a nuclear functional complex by analysing RNA bound to cytoplasmic and nuclear Ago2 in MCF-7 cells (GSE66665) (Supplementary Table 4). Surprisingly, the svtRNA1-2 fragment was enriched in both nuclear as well as in cytoplasmic Ago2 fractions. We used miR-21 [30] as a nuclear positive control (**Fig. S4B**).

Next, we wished to visualise endogenous vtRNA1-2 and svtRNA1-2 using Fluorescence *in situ* hybridisation (FISH). We designed probe 1 complementary to the middle region of vtRNA1-2 (the most unique region for vtRNAs in terms of sequence) for detection of full length vtRNA1-2. We also designed probe 2 complementary to 3’ region of vtRNA1-2 for detection of both full length vtRNA and svtRNA1-2 (**Fig. 2B**). First, we tested whether these probes specifically recognise vtRNA1-2 and not vtRNA1-1. Incubation of the probes with *in vitro* transcribed vtRNA1-2 and 1-1, followed by Northern blot shows that both probes are specific for vtRNA1-2 and do not hybridise to vtRNA1-1 (**Fig. 2B).** Next, we fluorescently labelled both probes and performed the FISH experiment in HEK293 WT and vtRNA KO cells. We observed that probe 1 gave a predominantly cytoplasmic signal, corresponding to full length vtRNA1-2, whilst in contrast, the signal for probe 2, corresponding to vtRNA1-2 and svtRNA1-2, was more nuclear. This was the case for FISH with incubation with single probes (1 or 2), or with double probe incubation (1+2). The quantification of fluorescent FISH signals confirms that the probe 2 signal is significantly more nuclear, whilst the probe 2 signal is more cytoplasmic. Using probes 1 and 2 in vtRNA KO cells did not result in positive signals, further confirming specificity of the FISH experiment (**Fig. 2C and D).** Closer inspection of the FISH signals also revealed the formation of probe 2 clusters, which were not visible with probe 1 (**Fig. S5A).** These data suggest that endogenous vtRNA1-2 resides more in the cytoplasm, whilst svtRNA1-2 is localised predominantly in nuclear clusters. Next, we transfected HEK239T cells with fluorescently labelled single- and double-stranded svtRNA1-2 and followed its cellular localisation at 4, 6 and 24 hours post-transfection. We observed mostly nuclear localisation of single-stranded and double-stranded svtRNA1-2 at all time points tested (**Fig. S5B and C**). We noted that transfected svtRNA were forming clusters, similarly to transfected tsRNA [9], but not siRNA. These foci could potentially suggest functional aggregates and possibly are an exaggeration of the clusters formed by endogenous svtRNA1-2, as the synthetic svtRNA1-2 is added to the cells in an excess.

Taken together, this data shows that svtRNA1-2 fragments bind to Ago2 in both cellular compartments and can be detected in the nucleus, suggesting that it might play a role in nuclear gene silencing.

### svtRNA1-2 targets selected genes at the nascent RNA level

The Dicer dependency and association of svtRNA1-2 with Ago2, along with its nuclear localisation, prompted us to investigate whether svtRNA1-2 could regulate gene expression on a nascent RNA level, similar to tsRNA. We employed ChrRNA-seq to characterise the nascent transcriptome of HEK293T cells in WT, Ago2 KD and Dicer KD conditions [9] (**Fig. 3A** and Supplementary Table 4). Differential expression analysis revealed that Ago2 and Dicer KD had a broad effect on the transcriptome, with 1346 genes down-regulated and 2568 up-regulated in Ago2 KD, and 1052 genes down-regulated and 2391 up-regulated in Dicer KD (log2FC > 1/ log2FC < −1 and p-adj < 0.001). The up-regulated genes in Ago2 and Dicer KD conditions included mostly protein-coding genes and long non-coding RNA (**Fig. 3B and C** and Supplementary Table 2). In order to identify the role of svtRNA1-2 in gene silencing, we analysed the intersection of up-regulated protein-coding genes in both Ago2 and Dicer KD conditions (1272 genes), as we showed that svtRNA1-2 is Dicer-dependent and binds to Ago2 (**Fig. 3D**). We employed the well-known and widely used MiRanda software [31] for *in silico* prediction of svtRNA1-2 gene targets through sequence complementarity, similar to miRNA binding sites, with restricted seed length and binding energy (see Methods). Interestingly, we found that the highest number of predicted gene targets are unique for svtRNA1-2 (227 genes, Supplementary Table 2), when compared to svtRNA1-1 (78 genes) or svtRNA1-3 (83 genes) (**Fig. 3D**). We used microRNA-4487 (miR-5587) as a negative control, as it has a sequence similar to svtRNA (**Fig. 3D** and **Fig. S4C**). Gene ontology enrichment analysis of the predicted svtRNA1-2 target genes revealed pathways involved in cell migration, motility, and regulation of proliferation and growth (**Fig. 3E**). Indeed, transmembrane receptor tyrosine-protein kinase ERBB4 was among the top 5 predicted target genes with the highest log2 fold change (Supplementary Table 2). To investigate this further, we mapped the predicted binding sites along the transcript lengths and found a distinct density pattern for svtRNA1-2 distributed along the body of the genes. In contrast, the other svtRNA fragments were mapped mostly to the beginning of the transcripts, resembling miRNA mediated gene activation. It would be interesting to test whether vtRNA1-1 and 1-3 could lead to gene activation. (**Fig. 3F**). These data suggest different roles for svtRNA1-2 and 1-1/1-3. Furthermore, we observed that svtRNA1-2 mapped to intronic regions with the highest occurrence, similarly to tsRNA, suggesting a potential role of svtRNA1-2 in NRS (**Fig. 3G**). In summary, we identified genes that are up-regulated upon Dicer and AGO2 KD on a nascent level and are *in silico* predicted to be targeted by svtRNA1-2 mostly within introns.

**Figure 3.**
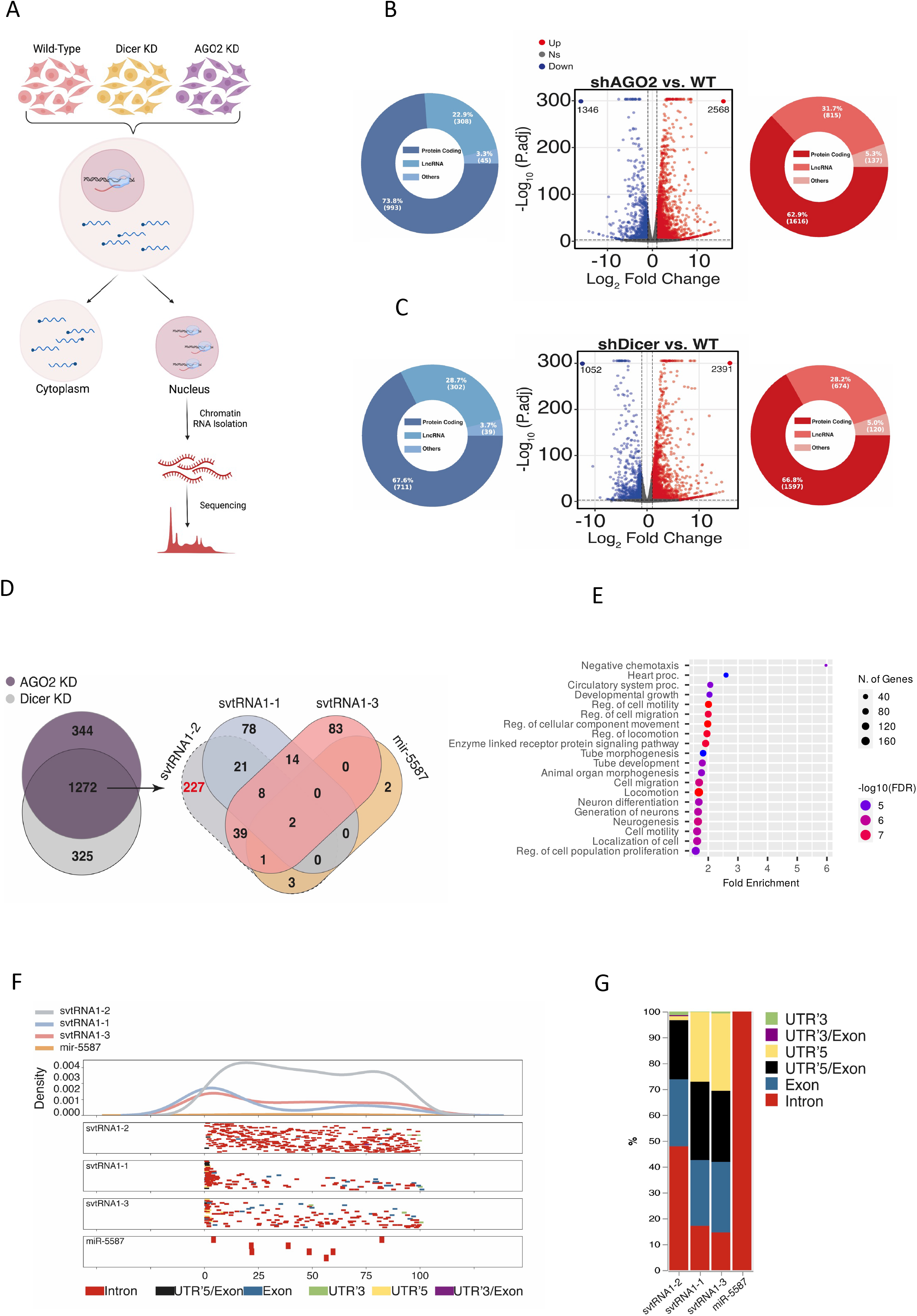
svtRNA1-2 targets selected genes at the nascent RNA level. **A)** Schematic representation of ChrRNA-seq method. **B)** Volcano plots of differentially expressed genes in AGO2 KD conditions. The red points show up-regulated genes and blue points the down-regulated genes with log2FC > 1/log2FC < −1 and p-adj < 0.001 respectively. Pie-charts show different gene types of up (red) or down (blue) regulated genes. **C**) As in B using Dicer KD samples. **D)** Venn diagram shows an overlap of up-regulated protein-coding genes from Dicer and AGO2 KD conditions (left). Venn diagram shows the number of predicted target genes for svtRNA1-1/2/3 and miR-5587 (right). svtRNA1-1/3 and miR-5587 were used as controls as their sequence similarity has the highest score with svtRNA1-2. **E)** GO analysis of vtRNA1-2 predicted targets upregulated in AGO2 and Dicer KD samples. **F)** Metagene distribution of predicted svtRNA binding sites mapped along normalised target transcripts length and their corresponding gene features. **G)** svtRNA binding site counts normalised to the length of a feature displayed in overall percentage.

### svtRNA1-2 knockout leads to defects in cellular proliferation

To investigate the molecular function of svtRNA1-2 further, we employed an RNA-guided CRISPR Cas12a system targeting vtRNA1-2 with single guide RNAs (sgRNA) in HEK239T cells (**Fig. S6A and B**). We successfully generated individual vtRNA1-2 knock-out (KO) clones and confirmed partial deletion of vtRNA1-2 by sequencing and PCR (KO results in a shorter genomic fragment). As vault loci show relatively high sequence similarity, we tested whether the other vtRNA loci remain intact in vtRNA1-2 KO. Indeed, vtRNA1-1, 1-3 and 2-1 genomic loci were not affected by vtRNA1-2 deletion (**Fig. 4A**). Next, we wished to validate the svtRNA1-2 predicted targets. We performed a qRT-PCR using RNA isolated from WT and stable vtRNA1-2 KO cell lines to validate the expression levels of ERBB4, ADAM12, KCNH1, PLXNA4 (unique targets for svtRNA1-2) and PARP12 (a unique target for svtRNA1-1, used as negative control). We show that the expression of all svtRNA1-2 predicted targets was significantly increased at both the nascent and steady-state RNA level in vtRNA1-2 KO cells, whilst the expression of PARP12 remained unaffected (**Fig. 4B and C**).

**Figure 4.**
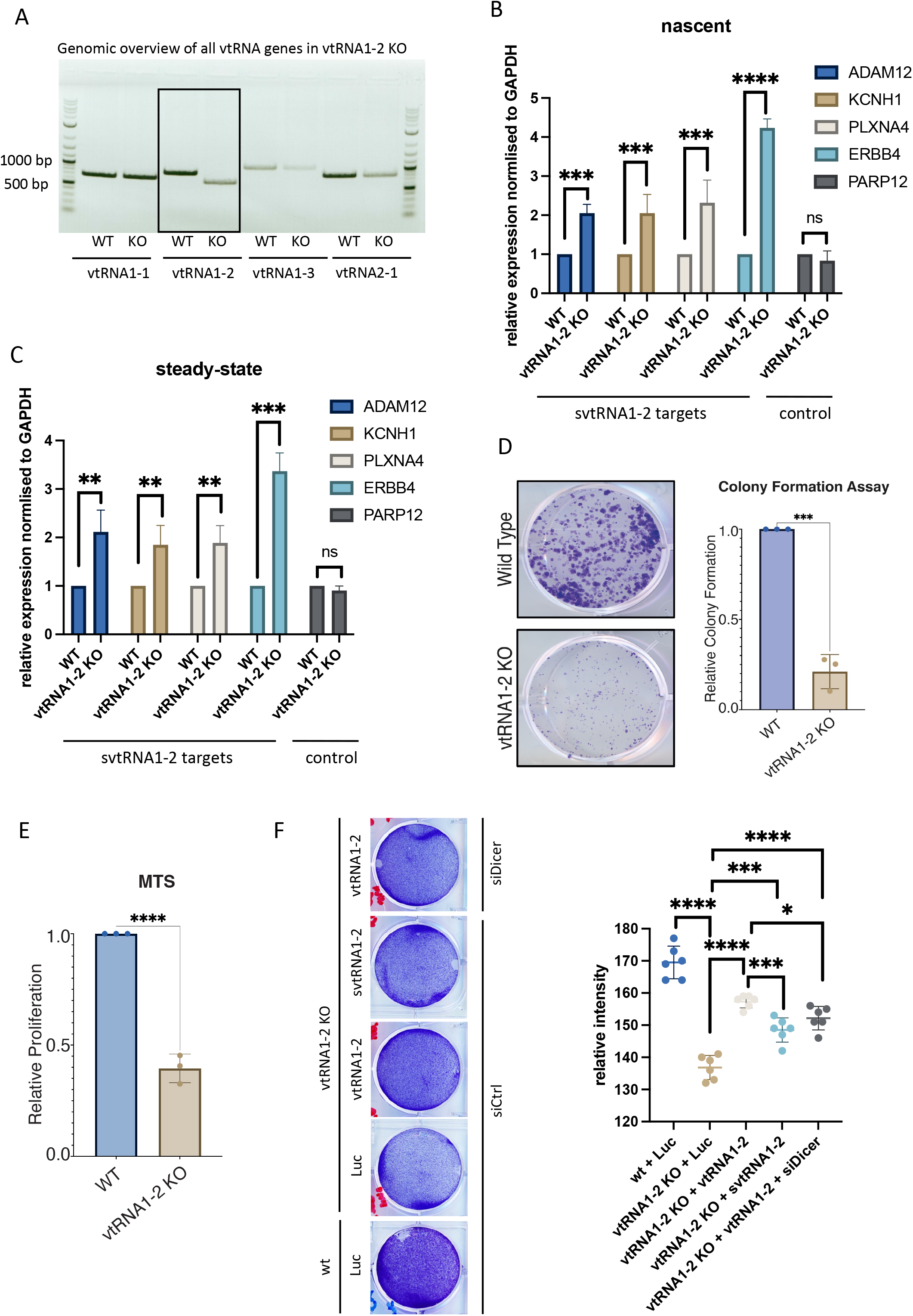
vtRNA1-2 Knock-Out leads to severe cellular phenotype. **A)** PCR analysis from purified genomic DNA from WT and vtRNA1-2 KO showing each vault RNA **B**) qRT-PCR quantification of nascent RNA levels of ADAM12, KCNH1, PLXNA4 and ERBB4 as predicted targets of svtRNA1-2 and PARP12 as a predicted target of svtRNA1-1 (used as negative control) in WT and svtRNA1-2 KO cells normalised to GAPDH. The data represent the mean fold changes from three independent experiments (****p< 0.0001; ***p< 0.001; n.s not significant). **C)** as in B, showing steady state RNA levels. **D)** Clonogenic assay of WT and vtRNA1-2 KO cells. The cells were stained and counted after 10 days of growing (*** P <0.001). **E)** MTS assay showing proliferation of WT and vtRNA1-2 KO cells after 7 days (****P < 0.0001). **F)** Left, representative image of proliferation assay of WT and vtRNA1-2 KO cells, transfected with plasmids expressing either control luciferase (Luc), full vtRNA1-2 or svtRNA1-2 in the presence of control siRNA (siCtrl) or siRNA targeting Dicer (siDicer). Right, quantification of relative intensity of a signal in rescue proliferation assay (****p< 0.0001; ***p< 0.001; *p< 0.005).

Interestingly, we observed impaired cellular proliferation in the vtRNA1-2 KO cells, when compared to WT cells, as shown by the clonogenic survival assay and MTS proliferation assay (**Fig. 4D** and **E**). The decreased cell viability and reduced proliferation of vtRNA1-2 KO cells, suggest that vtRNA1-2 plays an important role in normal cellular function. To validate this further investigate this phenotype, we performed a rescue experiment with plasmids expressing vtRNA1-2, svtRNA1-2 and luciferase plasmids (as a negative control). We show that vtRNA1-2 plasmid (providing both full and small vtRNA1-2) leads to significant rescue of proliferation phenotype. Furthermore, svtRNA1-2 alone also leads to partial but significant rescue, as well as vtRNA1-2 alone (vtRNA1-2 in Dicer KD in vtRNA1-2 KO cells). These data suggest that both vtRNA and svtRNA1-2 are relevant to vtRNA1-2 KO proliferation phenotype **(Fig. 4F).**

### Deletion of vtRNA1-2 alters the expression of genes associated with cellular membrane function

To further explore the role of svtRNA1-2 in nascent RNA gene silencing, we isolated chromatin-associated RNA from HEK293T WT and vtRNA1-2 KO cells and performed ChrRNA-seq (**Fig. 5A**). The purity of the fractions was confirmed by western blotting using H3 as a chromatin marker and Tubulin as a cytoplasmic marker (**Fig. S7A**). A principal-component analysis outlined clear differences between the two conditions and tight clustering of the replicates (**Fig. S7B and C**). Differential expression analysis revealed 547 down-regulated and 944 up-regulated genes (mostly protein-coding genes) in vtRNA1-2 KO cells (**Fig. 5B and Fig. S7D and E**). Closer inspection showed that ERBB4 was among the upregulated genes in vtRNA1-2 KO, providing further validation for svtRNA1-2 mediated downregulation of ERBB4 (**Fig. 5B** and **C**).

**Figure 5.**
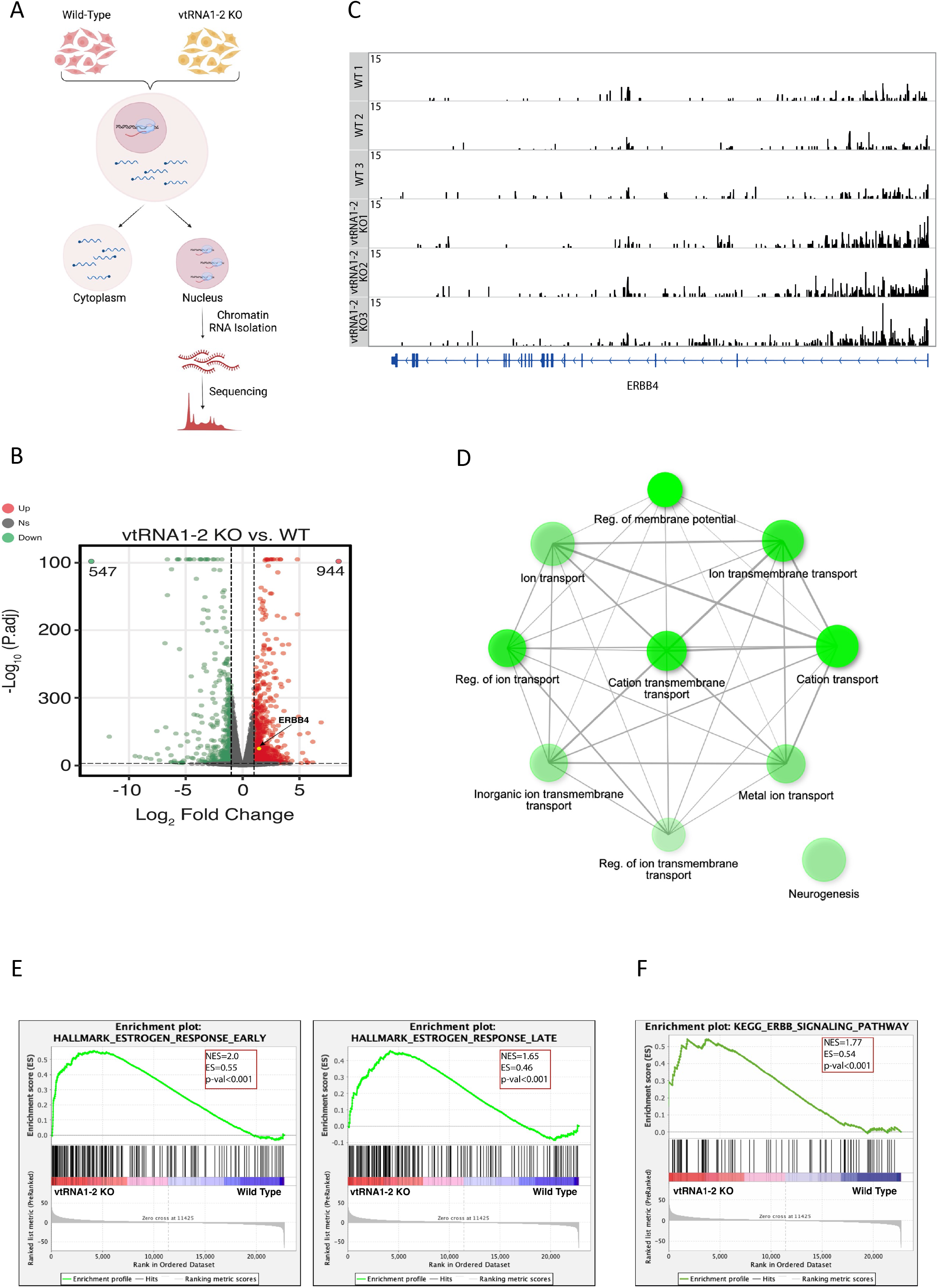
Deletion of vtRNA1-2 alters the expression of genes associated with cellular membrane function. **A)** Schematic representation of ChrRNA-seq method. **B)** IGV tracks of ChrRNA-seq from WT and vtRNA1-2 KO cells displaying the ERBB4 reads. The bottom blue line indicates the composition of ERBB4 gene. **C)** Volcano plots of differentially expressed genes in vtRNA1-2 KO and WT conditions. The red points show up-regulated genes and green points the down-regulated genes with log2FC > 1/log2FC < −1 and p-adj < 0.001 respectively. The yellow point shows ERBB4. **D)** GO analysis of up-regulated genes in vtRNA1-2 KO when compared to WT. **E)** Gene Set Enrichment Analysis (GSEA) with gene set from vtRNA1-2 KO cells using Hallmark gene sets in MSigDB. Genes are ranked by Wald statistic values in a ChrRNA-seq experiment (KO vs. WT). A positive enrichment score indicates higher expression after vtRNA1-2 KO. **F)** GSEA with gene set from vtRNA1-2 KO cells using KEGG pathways.

Ontology enrichment analysis of genes upregulated in vtRNA1-2 KO cells revealed an association of protein-coding genes with biological processes such as regulation of membrane transport and potential (**Fig. 5D** and **S8A**), while down-regulated genes were not enriched in these categories (**Fig. S8B**). Interestingly, enrichment analysis that includes both up and down-regulated genes revealed a significant association of differentially expressed genes upon vtRNA1-2 KO with early and late estrogen response, potentially, at least partially, due to deregulated ERBB4 (**Fig. 5E**). Finally, KEGG pathway analysis confirmed altered ERBB4 signalling pathway (**Fig. 5F**).

This data suggests that vtRNA1-2 is involved, either directly and/or indirectly, in the regulation of membrane proteins through the silencing function of svtRNA1-2.

### SvtRNA1-2 target genes are associated with cellular membrane function

Next, we used MiRanda software to predict target genes for svtRNA fragments in vtRNA1-2 KO up-regulated protein-coding genes. As expected, we identified most of the unique target genes for svtRNA1-2 (**Fig. S9A**). Closer inspection of MiRanda predictions revealed 139 target genes and 205 target sites for svtRNA1-2 (**Fig. S9B**). Comparing MiRanda with another target site prediction software PITA, we observed a good overlap of unique target genes, however more stringent MiRanda settings resulted in more predicted genes (**Fig. S9C** and Supplementary Table 5 and 6). The MiRanda software is based on dynamic programming local alignment between the query sequence and the reference sequence. High-scoring alignments estimate the thermodynamic stability of RNA duplexes based on these alignments. PITA tool assesses the accessibility of the target site. It evaluates the free energy gained from miRNA–mRNA pair formation and the energy cost of making the target accessible to the miRNA and computes the difference between these two parameters. Therefore, for further analysis, we used targets identified by MiRanda. Metagene alignment showed an even distribution of svtRNA1-2 target sites, along the gene length, whilst svtRNA1-1 and 1-3 target sites were accumulated at the beginning of the genes, similar to what we observed in **Fig. 3F** (**Fig. S9D**). Finally, svtRNA1-2 target sites were predominantly mapped to the intronic regions. These results are consistent with the previous predictions (**Fig. S9E**).

Focusing on unique svtRNA1-2 target genes, we analysed whether previously predicted svtRNA1-2 target genes that were upregulated upon Dicer and Ago2 knockdown and predicted to be targeted by svtRNA1-2 were also upregulated in vtRNA1-2 KO cells. We found 29 common genes including ERBB4 (**Fig. 6A**, **Fig. S9F** and Supplementary Table 3). In order to analyse whether this is statistically significant we performed further analysis as followed: first we used the 584 upregulated protein coding genes in the vtRNA1-2 KO sample and ran a random sub-set of protein coding genes 1000 times and overlapped this random subset with the 584 upregulated genes. This resulted in Min. : 0.000 1st Qu.: 7.000 Median : 9.000 Mean : 8.936 3rd Qu.:11.000 Max. :19.000, suggesting that the intersection of 29 upregulated vtRNA1-2 KO genes with the 227 upregulated genes in Dicer/Ago2 KD samples is indeed statistically significant. Finally, overall probability statistics revealed the following: Set1: 227, Set2: 584, Overlap: 29; total number of genes: 14759; Representation factor: 3.2; p < 2.510e-08. It should be noted that a representation factor > 1 indicates more overlap than expected of two independent groups, a representation factor < 1 indicates less overlap than expected, and a representation factor of 1 indicates that the two groups by the number of genes expected for independent groups of genes.

**Figure 6.**
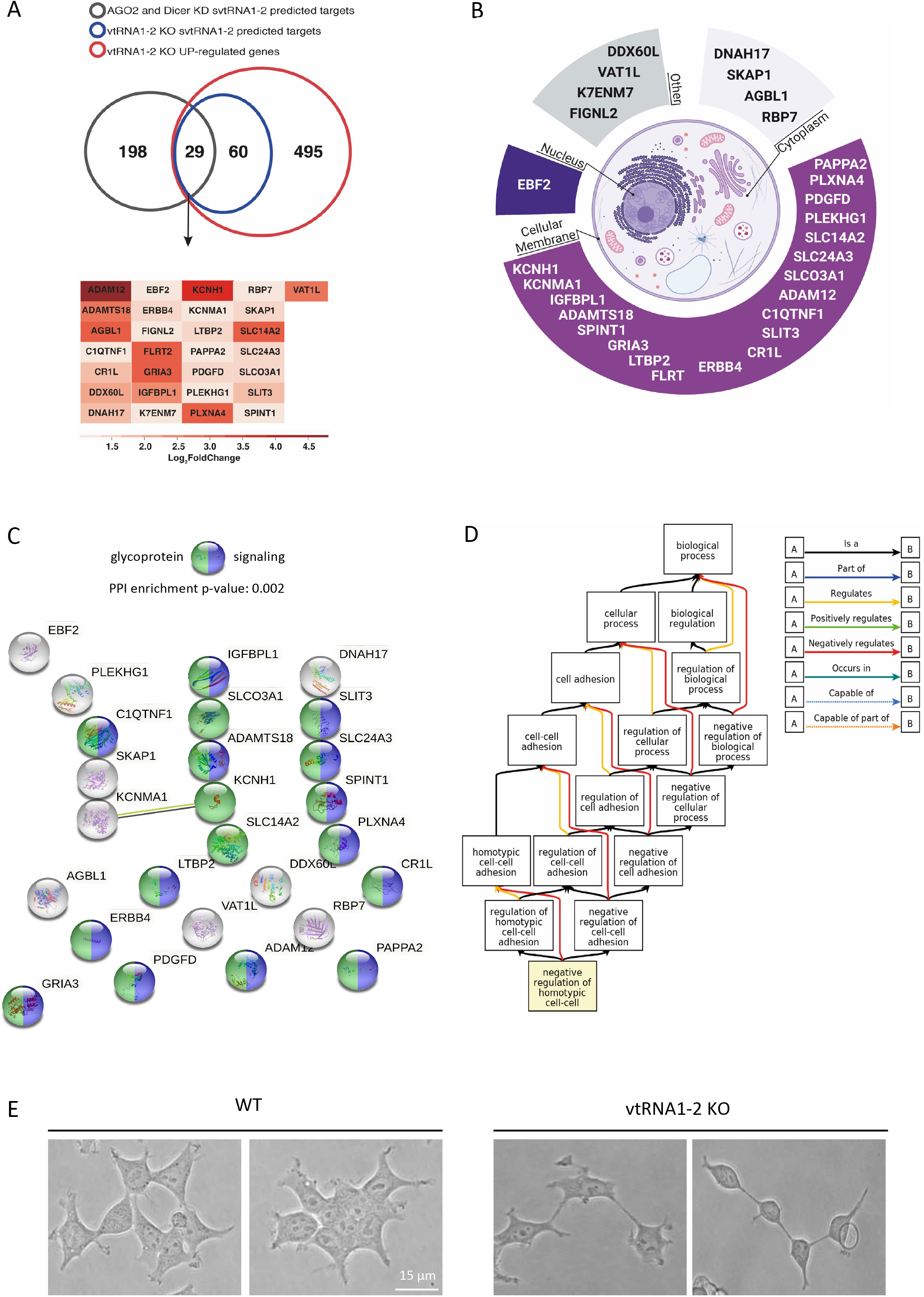
svtRNA1-2 target genes are associated with regulation of cell adhesion. **A)** Venn diagram showing the overlap between up-regulated protein-coding genes from vtRNA1-2 KO (compared to WT), vtRNA1-2 KO svtRNA1-2 predicted targets and svtRNA1-2 predicted targets from up-regulated genes in AGO2 and Dicer KD cells (top). Tables showing names of 29 svtRNA1-2 target genes. Heat map is showing fold of expression change for each target genes in vtRNA1-2 KO cells. **B)** Pie chart showing cellular localisation of 29 svtRNA1-2 target genes. **C)** STRING analysis showing interactions among 29 svtRNA1-2 target genes. Proteins involved in signalling are marked in blue, and glycoproteins are marked in green. **D)** Pathways analysis showing association of svtRNA1-2 target genes with negative regulation of cell adhesion. **E)** Representative examples of bright field microscopy showing differences in cell adhesion phenotype in WT and vtRNA1-2 KO cells.

Interestingly, proteins encoded by these genes were mostly localised in the cellular membrane (**Fig. 6B**). Indeed, STRING [32] analysis identified 14 out of 29 to be involved in signalling and 17 out of 29 to be glycoproteins (**Fig. 6C**). Glycoproteins are membrane proteins that play a key role in cell migration, proliferation, and adhesion [33]. Indeed, complex pathway analysis revealed that svtRNA1-2 target genes are involved in cell adhesion regulation (**Fig. 6D**). To validate the result, we seeded an equal number of WT and vtRNA1-2 KO cells and monitored their adhesion abilities. We observed elongated vtRNA1-2 KO cells with reduced cell to cell contact, most likely as potential consequence of negatively regulated cell-to-cell adhesion (**Fig. 6E**).

Overall, these data demonstrate that vtRNA1-2 is involved in the regulation of nascent RNA of protein-coding genes that are associated with cell membrane physiology.

### svtRNA1-2 targets intronic regions through a seed-plus-complementary-paired region and target loop protrusion architecture

Analysis of the MiRanda output revealed that svtRNA1-2 hybridises to the predicted targets in a miRNA-like fashion. Thermodynamic asymmetry between the two ends of the svtRNA1-2-target duplex, results in near-perfect complementarity at the 5’seed region and the 3’supplementary base pairing features depicted in the predicted targets (Supplementary Table 5). Interestingly, the analysis of svtRNA1-2-target hybridization features also revealed: 1) the nearly perfect base pairing extends over the 5’ seed region (2-8) up to nucleotide 18 with wobble base pairs and 2) the presence of a 5-nt loop in the target RNA for 179 out of 205 predicted targets. Moreover, the 5-nt loop shows a conserved sequence (CAUCA/CAUUA) within the 179 predicted target genes (Supplementary Table 5).

In order to determine the relationship between the svtRNA1-2 sequence and target hybridization, a comparative analysis of svtRNA1-2 WT and scrambled svtRNA1-2 sequences (5’shuffled, 3’shuffled, swapped) was performed on the group of the up-regulated genes in vtRNA1-2 KO (**Fig.7A** 5’ seed sequence is highlighted in orange, 3’ complementary sequence is highlighted in blue, shuffled bases are depicted in red; svtRNA1-1 was used as negative control). To prevent a biasing artefact, we kept the base composition of shuffled svtRNA sequences similar (**Fig. S10A**). All svtRNA sequences were run through MiRanda software against up-regulated genes identified in vtRNA1-2 KO to obtain their target predictions. Mapping of 5’ shuffled svtRNA1-2 sequence resulted in complete loss of targets when compared to the svtRNA1-2 WT sequence, whilst 3’ shuffled svtRNA1-2 mapped to only 3 targets.

**Figure 7.**
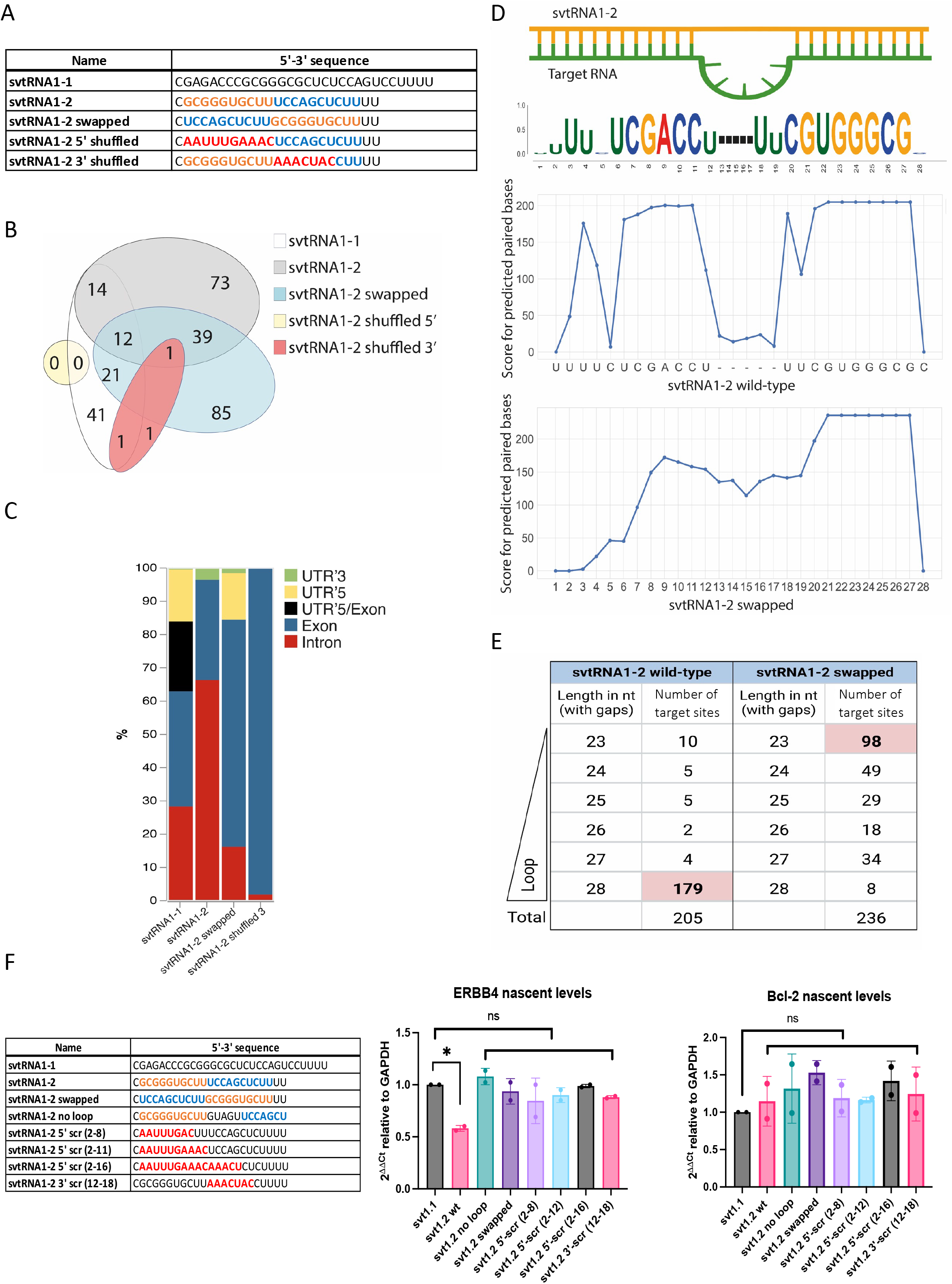
svtRNA1-2 targets intronic regions through a seed-plus-complementary-paired region and target loop protrusion architecture. **A)** Sequence representation of svtRNA1-1, svtRNA1-2 WT, svtRNA1-2 swapped, svtRNA1-2 shuffled 5’ and svtRNA1-2 shuffled 3 ‘. **B)** Venn diagram showing an overlap of target genes predicted for each sequence. **C)** Stacked bar plot showing predicted svtRNA binding sites mapped to different gene features with normalised counts to the length of a feature._Displayed in overall percentage. **D)** Sequence logo of significantly enriched bases in svtRNA1-2 predicted target sites (top). Line plot representing scores for predicted paired bases of svtRNA1-2 to their target sites (middle). Line plot representing scores for predicted paired bases of swapped svtRNA1-2 to their target sites (bottom). **E)** Table showing correlation analysis of the loop/no loop and number of target sites. 23 nt represents alignment with no loop, 28 nt contains a 5 nt long gap across the alignment, loop (the longest loop observed). F) Left, table of sequences used for transient transfection of MCF-7 cells. Right, bar chart showing nascent RNA levels of ERBB4 (svtRNA1-2 target gene) and Bcl2 (control gene) measured by qRT-PCR. Samples were isolated 24 hours after trasfection. The values were normalised to the GAPDH. The data represent the mean fold changes from three independent experiments (*P < 0.05; n.s not significant).

The swapped svtRNA1-2 mapped to 158 targets which partially overlap with the svtRNA1-2 WT predicted targets (39 out of 138) (**Fig.7 B**). However, unlike the svtRNA1-2 wt, the swapped svtRNA1-2 sequence predominately hybridised to the exons of the target genes (**Fig. 7 C and Fig. S10B**), indicating that the predicted swapped svtRNA1-2 targets have different binding sites along the nascent RNA target when compared to svtRNA1-2 WT sequence. Of note, the 5-nt loop structure was not detected within the group of targets predicted for the swapped svtRNA1-2 (**Fig. 7 D and E**). The majority (147 out of 236) of the predicted swapped svtRNA1-2 target sites showed a length of 23 or 24 nucleotides that corresponds to the absence of the loop structure (**Fig. 7 E**).

To gain further insight into the molecular mechanism of svtRNA1-2-mediated NRS, several modified svtRNA1-2 molecules were transfected into the human breast cancer cell line MCF-7, and the expression level of nascent ERBB4 RNA was assessed by RT-qPCR, 24 hour post transfection (**Fig.7 F**).

Specifically, we employed the single stranded svtRNA1-2 molecules: 5’ scrambled at nucleotides 2-8, 2-12 and 2-16, 3’ scrambled at nucleotides 12-18, swapped at 5’/3’ ends, and svtRNA1-2 bearing perfect complementarity with the target gene (svtRNA1-2 no loop). Interestingly, the nascent RNA level of ERBB4 was significantly reduced by svtRNA1-2 WT transfection, but was not altered in presence of svtRNA1-2 with no loop (perfectly complementary), indicating that the 5-nt loop on the target RNA is pivotal for NRS of ERBB4. Moreover, loss of silencing upon transfection of 5’ and 3’ scrambled svtRNA1-2 sequences indicates that complementarity of both the 5’ and 3’ regions is fundamental for effective NRS. The absence of silencing with the swapped svtRNA1-2 also suggests that silencing of ERBB4 is sequence and site specific. Finally, Bcl-2 nascent RNA levels, used as negative control, were not affected by svtRNA1-2 molecules.

These data demonstrate that svtRNA1-2 possess unique features that are required for specific intronic targeting for NRS.

## Discussion

In this work, we have identified Dicer-dependent small RNA fragments derived from vtRNA1-2 and vtRNA2-1. We also found vtRNA1-1 to be independent of Dicer processing in HEK293T cells, which is in line with previous observations by Rybak-Wolf *et al*. [24]. However, it is worth noting that there are also contrasting results that show vtRNA1-1 as a Dicer substrate [34, 35]. This discrepancy could be potentially explained by the different sensitivity of the experimental techniques and also perhaps by differing efficiencies of Dicer KD. Furthermore, vtRNA are also differentially expressed in various tissues [13, 22, 36, 37], so vtRNA1-1 might be processed by Dicer in other cell lines. The analysis of RNA bound to Argonaute proteins using two different experimental methods identified svtRNA fragments derived from vtRNA1-1/2 to be bound to Argonaute proteins, whilst svtRNA2-1 fragments were not detected. In contrast, recent work showed the binding of Ago2 to the 5’ vtRNA2-1 fragment by qRT-PCR in SH-SY5Y cells [38]. The potential explanation for this contrasting observation could be a differential expression of vault RNA in various tissues and pathological contexts. Genome-wide expression studies of small non-coding RNA, including svtRNA, in different cell types or tissues, would allow the identification of more detailed vtRNA expression patterns and could provide important insights into their biogenesis.

We found svtRNA1-2 to be associated with Ago2 not only in the cytoplasm but also in the nucleus. The nuclear localisation of svtRNA1-2 was confirmed by confocal microscopy using FISH and synthetic fluorescently labelled svtRNA1-2. As vtRNA share similar features with tRNA, we hypothesised that svtRNA1-2 could guide Ago2 in the nucleus to promote NRS [9]. Furthermore, Ago2 has previously been proposed to regulate gene expression in the nucleus [28].

In order to investigate the potential role of svtRNA1-2 in an NRS mechanism, we used chromatin-associated RNA sequencing datasets from Dicer and Ago2 KD cells and predicted svtRNA1-2 binding sites. We found that svtRNA1-2 mapped with the highest occupancy to the intronic regions in many genes that are involved in cell membrane physiology, including ERBB4. Interestingly, genomic deletion of vtRNA1-2 resulted in inhibition of cell growth and proliferation. Such dramatic defects in cell growth and proliferation were not seen previously in vtRNA1-1 and vtRNA1-3 KO cells [39], suggesting that vtRNA1-2 might play a role in different cellular processes. Analysis of nascent RNA isolated from HEK293T WT and vtRNA1-2 KO further confirms the role of svtRNA1-2 in NRS.

Well-established rules for functional miRNA-target pairing, classify target sites into 3 main categories; canonical sites that pair at both 5’ and 3’ends, seed sites (depending on minimal 5’end base pairing), and 3’compensatory sites characterised by weak 5’base pairing and strong compensatory pairing to the 3’end [40]. To assess the NRS requirements of the svtRNA1-2 molecule, we analysed svtRNA1-2 WT pairing conditions between svtRNA1-2 and the predicted targets. A large set of predicted targets for svtRNA1-2 WT showed perfect 5’seed complementarity with extended wobble base pairs at the 3’end. Nucleotide conservation analysis of the predicted targets (**Fig. 7D**), highlighted the presence of a GC-enriched segment at the 5’end that might have additional roles such as the 5’ Terminal Oligo Guanine (TOG motif) described as a characteristic feature of some 5’ tsRNA [41]. We cannot exclude that svtRNA1-2 might form intermolecular RNA G-quadruplex structures and gain further functions in translational inhibition, protein recognition, or cellular localization. In addition, svtRNA1-2 central (10-11-12) positions predominantly follow the base pairs scheme Wobble-WC-Wobble. It should be noted that most of the predicted targets have an unpaired loop of 5 nucleotides opposite to positions 11 and 12 of the svtRNA1-2, with a common sequence 5’-ACUAC or AUUAC. It has been demonstrated that Ago2 can accommodate a central loop present on the target gene, without loss of binding affinity [42]. Furthermore, we hypothesised that the target loop might be recognised as a consensus motif by some proteins. To further evaluate the minimal pairing requirements between svtRNA1-2 and the targets, we designed two svtRNA1-2 molecules with a shuffled sequence at either the 5’end or 3’supplementary region (shuffled svtRNA1-2 5’ and shuffled svtRNA1-2 3’, respectively).

Interestingly, the svtRNA1-2 shuffled 5’ designed to follow the rules of thermodynamic asymmetry, present in large numbers of miRNA molecules, did not have any common target genes with svtRNA1-2 WT (**Fig. 7B**). Loss of common targets could be connected to higher free energy values of shuffled svtRNA1-2 5’ with respect to the WT svtRNA1-2. Shuffled svtRNA1-2 at the 3’end showed similar results to the svtRNA1-2 shuffled at the 5’end, resulting in no common targets with svtRNA1-2 WT (**Fig. 7B**). This data demonstrates that complementarity at both the 5’ and 3’ends of the svtRNA1-2 is crucial for the identification of target genes. Since we reasoned that the complete loss of common targets between WT and 5’ and 3’ shuffled svtRNA1-2 might be partially due to the MiRanda free energy cut-off, we designed a svtRNA1-2 molecule with a swapped 5’ and 3’ seed sequence. In this way, we preserved not only the nucleotide composition but also the sequence identity, while following the rules for the design of efficient miRNA molecules. From the analysis of the target predictions, we observed some increase in the putative targets and a dramatic change in the target sequence complementarity localisation. The swapped svtRNA1-2 hybridises within the exons of the predicted genes, in contrast to the svtRNA1-2 WT which hybridises within introns via loop protrusion. This data sheds light on the importance of the G-rich region at the 5’end for the svtRNA1-2-mediated NRS.

## Conclusion

In conclusion, in this work, we uncovered a Dicer-dependent small RNA fragment derived from vault RNA1-2. We show that svtRNA1-2 mediates nuclear gene silencing by targeting the introns of prote-incoding genes that are associated with cell membrane function (**Fig. S10C**). Deletion of vtRNA1-2 results in a deregulated proliferation phenotype, which underpins the biological significance of vtRNA1-2. Finally, we show that svtRNA1-2 employs double complementarity with the formation of a loop protrusion within its target sequences, which is required for intronic binding. Although we cannot yet fully appreciate how the loop protrusion might be recognised and used for nascent RNA degradation, we demonstrate a novel biological role for vtRNA1-2 and its potential for future RNA-based cancer therapies.

## Materials and Methods

### Small RNA analysis

FASTQ files were trimmed to remove the 3’ adapter sequence (GATCGGAAGAGCACACGTCTGAACTCCAGTCACCGATGTATCTCGTATGCCGTCTTCTGCTTG) and filtered based on the length of the read (15 to 30) using Cutadapt (v.1.8.3)[43]. Following data pre-processing reads were then aligned with STAR aligner (v.2.5.3a)[44] to the human genome (hg38) downloaded from the GENCODE (v.34) with basic human genome annotation. Custom small non-coding RNAs annotation file was built by merging annotation of miRNA from miRbase (v.22.1)[45], tRNA from GENCODE (v.34) and snoRNA, snRNA, miscRNA, scaRNA, rRNA and vault RNA from GENCODE (v34) (hg38) basic genome annotation. Reads overlapping the custom annotation were quantified using featureCounts (v.1.6.2)[46]. DEGs were identified using the DESeq2[47] with an adjusted p-value < 0.05. The vault RNA fragments were identified using bedtools (v.2.29.2)[48] genomecov with the following settings –dz and –scale, which calculates the read coverage along the genome.

### 6xHis-Dicer purification

Low passage *Spodoptera frugiperda* (Sf9) cells were seeded at 0.5×10^^6^ cells/mL and grown in suspension at 110 rpm at 27°C. On the next day cells reached density of 1.5-2.5 ×10^^6^ cells/mL and were infected with P1 baculovirus at MOI=1, generated with pFastBac1-hDicer (Addgene, #89144), and Bac-to-Bac expression system (Gibco). All steps of purification were carried out at 4°C. Cells were harvested 72 hours post infection by spinning at 800 x g for 15 min and lysed with 6x volume of lysis buffer (20 mM TrisHCl pH 8, 500 mM NaCl, 10% glycerol, 0.5% Triton-X, 1mM TCEP, 1x Complete protease inhibitor, 1x PMSF, 1x Leupeptin) and micrococcal nuclease (NEB), and rotated on a wheel at 25 rpm for 30 min. The lysate was then sonicated 4 times for 30 sec on/off at 15 μm, span at 13000 x g for 45 min and then filtered with a 0.2 μm filter. 250 μL of Ni-NTA agarose (Qiagen) for 50 mL of cell culture were equilibrated with wash buffer (20 mM TrisHCl pH 8, 500 mM NaCl, 5% glycerol, 1mM TCEP, 1x Complete protease inhibitor, 1x PMSF, 1x Leupeptin and 5 mM imidazole). Imidazole was added to the lysate to 5 mM and then incubated with pre-equilibrated Ni-NTA agarose on a tube roller for 1 hour. Agarose-lysate mixture was span down gently at 300 rpm and resin was let set in a gravity flow column (BioRad) and washed twice with wash buffer. 6xHis Dicer was then eluted with elution buffer with a gradient of imidazole (20 mM TrisHCl pH 8, 500 mM NaCl, 5% glycerol, 1mM TCEP, 1x Complete protease inhibitor, 1x PMSF, 1x Leupeptin, 50 mM-500 mM imidazole). Fractions were run on SDS-PAGE, stained with Coomassie blue or immunoblotted (abcam #14601). Active fractions were concentrated, and buffer exchanged with a 100 KDa cut-off concentrator (Vivaspin) and stored in storage buffer (50 mM TrisHCl pH 8, 50 mM NaCl, 5mM MgCl_2_, 0.2% (mg/ml) BSA, 2 mM TCEP, 20% Glycerol, 1x Protease inhibitor) at −80 °C.

### *In vitro* Dicer Cleavage assay

1 μg of T7 *in vitro* transcribed substrates (vtRNAs, tRNAs and control ncRNAs) were slow-cooled annealed by boiling at 95°C for 3 min and then let at room temperature until temperature equilibration. Substrates were then incubated with 2 μg of recombinant purified Dicer and 1 μl of Ribolock RNAse inhibitor in Dicer reaction buffer (50 mM TrisHCl pH 8, 300 mM NaCl, 20 mM HEPES, 5 mM MgCl_2_, 5% glycerol and 1x protease inhibitor), and incubated for 4 hours at 37°C. Aliquots were collected at time points by mixing with 1X TBE-Urea loading dye (Novex) and immediately frozen on dry ice. Samples were run on a 10% urea polyacrylamide gel in 1x TBE at 450 V and then transferred onto Hybond-N + nylon membrane (Cytiva) for 1 hour at 5V, followed by UV-crosslinking and then pre-hybridized for 30 min in ULTRAhyb-Oligo buffer (Thermo Fisher Scientific) at 42°C. Probes were radiolabelled with 32P-ATP by poly nucleotide kinase (PNK) (PerkinElmer PNK) for 1 hour at 37 °C, purified using a G-25 Sephadex column (GE Healthcare) and then incubated with membranes overnight. Membranes were then washed 2x with 0.1x SCC buffer and subjected to autoradiography.

### Fluorescence In Situ Hybridization (FISH) for vtRNA1-2 and svtRNA1-2 detection

HEK293T, HEK293T vt1-2 ko and HeLa cells were seeded on coverslip to reach 40% of confluence after 24 hours. Cells were rinsed twice with PBS and fixed with PFA 4% in PBS for 15 minutes at 37 degrees. Samples were firstly incubated twice with 1-methylimidazole solution (0.13M 1-methylimidazole, 300mM NaCl in DEPC H2O, pH 8) at room temperature for 10 minutes and then with freshly prepared EDC (1-ethyl-3-(3-dimethylaminopropyl) carbodiimide hydrochloride) solution (EDC 0.1 M in 1-methylimidazole solution at pH 8) in humidified chamber at room temperature for 1h. Subsequently, samples were incubated twice with 1% w/v Glycine in PBS for 5 minutes at room temperature and rinse twice with PBS. Permeabilization step has been performed incubating samples with 0.2% Triton in PBS for 10 minutes at room temperature and rinse twice with PBS. SSC blocking buffer solution (SSC 2x, BSA 2%, sssDNA 40 ng/μl, yeast tRNA 40ng//μl, RNAsin Plus 0.67U/ μl in DEPC H2O) has been used prior probes hybridization step. Samples were incubated in humidified chamber at 37 degrees for 1h. FISH probe buffer (SSC 2x, sssDNA 30ng/μl, yeast tRNA 30ng/μl, RNAsin Plus 1U/ μl and 100nM or 50nM of fluorescent label probes) was added to the samples and let hybridise at 95 degrees for 5 minutes and 4 degrees overnight. Post-hybridization washes (3 washes at 4 degrees 5 minutes each) were performed using different SSC buffer solutions in DEPC H2O (SSC 2x, BSA 1%; SSC 1x, BSA 1%; SSC 0.5x, BSA 1%) and rinse twice with PBS.

### Identification of small vault RNA in mouse tissue samples

To investigate the possible expression of murine small vault RNA, I used sRNA-seq data from various mouse tissues, including (spleen, liver, lungs, muscle, brain, and pancreas) (GSE119661, Table 1). The data were obtained in triplicates for most of the tissues and then duplicates for (Brain and Heart) and processed according to published methods Kern et al. 2020. Briefly, obtained raw reads were processed and aligned to mouse genome index (m39) using STAR (v2.7.3a), and after the alignment, I generated normalised coverage files in bigWig format using bamCoverage function from deeptools (v2.2.2) and visualised them in IGV.

### Ago1-4 PAR-CLIP analysis

The Ago1-4 PAR-CLIP datasets adapter sequences were trimmed using Cutadapt (v.1.8.3)[43]. Then the reads were mapped to the reference human genome (hg38) by Bowtie (v.1.1.2)[49], allowing for two alignment errors (mutation, insertion or deletion). For each read, only the best mapping was reported out of a maximum of 10 genomic matches. Any tag with over 10 genomic matches was discarded. After the conversion subtraction, reads that mapped to only one genomic location were retained for further analysis. The clusters of reads (Ago binding sites) were identified based on T-to-C conversions using PARalyzer (v.1.5)[50] detailed settings described in (Supplementary Table 4). Then, these files were visualised using Integrative Genome Viewer (IGV)[51, 52].

### Ago2/3 RIP-seq

The Ago2/3 RIP-seq datasets consisted of 1 replicate per Ago2/3 sample (HEK293T cell line). Next, the 3’ adapter sequence (TCTCGTATCGTATGCCGTCTTCTGCTTG) was trimmed using fastx-clipper from fastx toolkit (v.0.0.13.2) from Hannon Lab with a quality score of 33 and read length longer than 15 nt. The index of the human genome was built with bowtie-build (v.1.1.2)[49] and reads mapped to the reference human genome (hg38) using Bowtie (v.1.1.2)[49]. The Stranded-Coverage tool from GitHub (https://github.com/pmenzel/stranded-coverage), was used to calculate the read coverage for vtRNA in the genome with the following settings -f, -x, -n (normalisation of the coverage (Reads Per Million)) -s 1. Output files were visualised using IGV.

### Ago2 RIP-seq Nuclear/Cytoplasmic

The reads were trimmed using cutadapt with the following settings -e 0.05 -q 23 -m 15 -M 30 -a “A[53]” and aligned to hg38 using STAR aligner allowing maximum of 3 mismatches and retaining only uniquely mapped reads. Next bamCoverage function was used from deeptools[54] to calculate coverage files using –binSize 1 and –normalizeUsing RPKM into BigWig files which were next visualised by IGV.

### Dicer PAR-CLIP analysis

FASTQ files (three biological replicates) were trimmed to remove the 3’ adapter sequence (TCTCACGTCGTATGCCGTCTTCTGCTTG,TCTCCATTCGTATGCCGTCTTCTGCTTG,TCTCC CATCGTATGCCGTCTTCTGCTTG) and select reads longer than 15 bases in size using Cutadapt (v.1.8.3). The alignment was done using Bowtie (v.1.2.3) with the following settings –v 3 –a –m1 –best –strata. The clusters of reads (Dicer binding sites) were identified based on T-to-C conversions using PARalyzer (v.1.5) detailed settings described in (Supplementary Table 4). Then, these files were visualised using Integrative Genome Viewer (IGV).

### ChrRNA-seq Dicer and Ago2 KD

The adapter sequences (-A AGATCGGAAGAGCGTCGTGTAGGGAAAGAGTGT and -a AGATCGGAAGAGCACACGTCT) were trimming from datasets (2 replicates per condition) by Cutadapt (v.1.8.3), discarding processed reads shorter than 10 nt. The processed reads were then mapped to the human reference genome (hg38) using STAR aligner (v.2.5.3a). The basic genome annotation was downloaded from the GENCODE website. The genes were quantified using featureCounts (v.1.6.2). Any gene with less than 5 read counts per replicate in at least two conditions were removed, and the counts’ matrix was then normalised using DESeq2.

### Generation of svt1.2 knockout cell line

Cas12a-gRNA ribonucleoprotein complexes containing one sgRNA targeting the vault1-2 gene (TTTAGCTCAGCGGTTACTTCGAGTACA) were nucleofected in HEK293T using Neon Transfection System (Thermo Scientific). After 48 hours cells were single-cell sorted into 96-well plates and subsequently genotyped. Cells bearing homozygous deletion of vtRNA1-2 was confirmed by Sanger sequencing.

### MTS Proliferation assay

WT and vtRNA1-2 KO cells were seeded into 96 well plates at density of 100 cells per well and left to proliferate for 7 days. The MTS assay was carried out using the CellTiter 96^®^ AQueous One Solution Cell Proliferation Assay (MTS) (Promega) according to the manufacturer’s instructions, and absorbance was measured at 490nm.

### Colony Formation Assay

To assess the impact of knockout of vtRNA1-2 on cell survival, colony formation assays were carried out. WT and vtRNA1-2 KO cells were seeded onto 12 well plates at density of 1000 cells per well and left for 10 days until visible colonies formed. Plates were stained with Crystal Violet staining solution (0.5% crystal violet, 20% ethanol) for 30 minutesand then quantified using the ColonyArea plugin on ImageJ.

### Rescue experiment of the HEK293T vt1-2 ko cell line

HEK293T vt1-2 ko cells were seeded into 6 well plate to reach 40% confluence the next day. siRNA against Dicer (siDicer) or non-silencing control siRNA (siNC) at concentration of 100nM have been transfected using, Lipofectamine RNAiMAX reagent following the manufacturer’s instructions. 24 hours after transfection cells were transfected with either pH1shluc or pH1vt1-2 or pH1svt1-2 vectors using Lipofectamine 3000 reagent following the manufacturer’s instruction. After additional 24 hours cells were harvested and seeded into 6 well plate at density of 3×105 and left to proliferate for 3 days. Finally, plates were stained with Crystal Violet staining solution (0.5% crystal violet, 20% ethanol) for 30 minutes and then quantified using ImageJ.

### Confocal and bright light microscopy

HEK293T cells were grown on a poly-L-lys coverslips and transfected with or 100nM of single-stranded or 40nM of double-stranded 5’-Cy3-labelled svtRNA1-2 24h after seeding. Cells were fixed at specific time points (4, 6, 24h) by paraformaldehyde (4% in PBS) and subsequently imaged on an FV1200 Olympus laser scanning confocal microscope equipped with an argon laser and a He/Ne 633 nm laser, using 60x/NA1.4 oil immersion objective. Comparative immunofluorescence analyses were performed maintaining identical acquisition parameters. Nuclear foci were analysed with Cellprofile software and statistical analysis was performed using GraphPad Prism.

Bright light microscopy was performed using 40x lens.

### Chromatin associated RNA-sequencing sample preparation

Chromatin RNA samples for sequencing were prepared according to previous publications[9, 55] with some minor modifications. Briefly, WT and vtRNA1-2 KO cells were harvested from 15 cm dishes and lysed in HLB+N buffer (10mM Tris-HCl (pH 7.5), 10 mM NaCl, 2.5 mM MgCl2, 0.5% NP-40), and underlaid with HLB+NS buffer (10 mM Tris-HCl (pH 7.5), 10 mM NaCl, 2.5 mM MgCl2, 0.5% NP-40, 10% sucrose), before centrifugation at 420g for 5 minutes. Nuclear pellets were then resuspended in NUN1 buffer (20 mM Tris-HCl (pH 7.9), 75 mM NaCl, 0.5 mM EDTA, 50% glycerol). Chromatin was extracted by addition of NUN2 buffer (20 mM HEPES-KOH (pH 7.6), 300 mM NaCl, 0.2 mM EDTA, 7.5 mM MgCl2, 1% NP-40, 1M urea) on ice for 15 minutes with vortexing. Samples were centrifuged at 17,000g for 10 minutes and chromatin pellets collected. Chromatin was digested by the addition of Proteinase K and Turbo DNase (Thermo Fisher). RNA was extracted from the chromatin samples using TRIzol (Invitrogen) and chloroform extraction, followed by the Monarch^®^ Total RNA Miniprep Kit (NEB) according to the manufacturer’s instructions and eluted in water.

### Western blotting

4x Laemmli buffer (0.2 M Tris–HCl, 8% (w/v) SDS, 40% glycerol, 20% (v/v) β-mercaptoethanol, 0.005% bromophenol blue) was used to treat the whole cell and chromatin extracts following 95°C incubation for 5 min and sonication. Samples were separated on mini-PROTEAN^®^ TGX™ gels (Bio-Rad Laboratories) followed by transfer onto nitrocellulose membrane (Protran, GE Healthcare) and probed with Tubulin (55kDa) and H3 (15kDa) antibodies.

### svtRNA1-2 transfection

MCF-7 cells were seeded to reach 70% of confluency at the time of transfection. 5’-phosphorylated single-stranded vtRNA1-2 at concentration of 100nM have been transfected using, Lipofectamine RNAiMAX reagent following the manufacturer’s instructions. 24 hours after transfection cells were harvested and RNA were extracted using TRIzol LS reagent.

### qRT-qPCR

1 μg of total RNA were reverse transcribed with gene specific primers using SuperScript III following the manufacturer’s instructions.

1:10 of cDNA were subsequently analysed using SYBR green-based qPCR assay and following the manufacturer’s instructions. Quantification of nascent RNA expression level of ERBB4 and BCl-2 genes was normalised to GAPDH RNA nascent expression level using the 2–ΔΔCt method. All RT-qPCR experiments were performed using three biological replicates.

### Primer sequences

List of all primers used in this study is available in Supplementary table 7.

### Plasmid construction for rescue experiment

DNA inserts bearing the human H1 promoter sequence (in bold) upstream either short hairpin luciferase (shluc) or vt1-2 (vt1-2) or small vt1-2 (svt1-2) sequences (underlined) have been obtained by gBlock gene fragments synthesis (IDT) and subsequently cloned into a plasmid backbone via Gibson assembly protocol (New England Biolabs) following manufacturer’s instructions. Recombinant plasmid sequences have been verified by Sanger sequencing.

(H1shluc)

GAACGCTGACGTCATCAACCCGCTCCAAGGAATCGCGGGCCCAGTGTCACTAGGCGGGAACACCCAGCGCGCGTGCGCCCTGGCAGGAAGATGGCTGTGAGGGACAGGGGAGTGGCGCCCTGCAATATTTGCATGTCGCTATGTGTTCTGGGAAATCACCATAAACGTGAAATGTCTTTGGATTTGGGAATCTTATAAGTTCTGTATGAGACCACAGATCCCCGGATTCCAATTCAGCGGGAGCCACCTGATGAAGCTTGACGGGTGGCTCTCGCTGAGTTGGAATCCATTTTT;

(H1vt1-2)

GAACGCTGACGTCATCAACCCGCTCCAAGGAATCGCGGGCCCAGTGTCACTAGGCGGGAACACCCAGCGCGCGTGCGCCCTGGCAGGAAGATGGCTGTGAGGGACAGGGGAGTGGCGCCCTGCAATATTTGCATGTCGCTATGTGTTCTGGGAAATCACCATAAACGTGAAATGTCTTTGGATTTGGGAATCTTATAAGTTCTGTATGAGACCACAGATCCCCGGGCTGGCTTTAGCTCAGCGGTTACTTCGAGTACATTGTAACCACCTCTCTGGGTGGTTCGAGACCCGCGGGTGCTTTCCAGCTCTTTTT;

(H1svt1-2)

GAACGCTGACGTCATCAACCCGCTCCAAGGAATCGCGGGCCCAGTGTCACTAGGCGGGAACACCCAGCGCGCGTGCGCCCTGGCAGGAAGATGGCTGTGAGGGACAGGGGAGTGGCGCCCTGCAATATTTGCATGTCGCTATGTGTTCTGGGAAATCACCATAAACGTGAAATGTCTTTGGATTTGGGAATCTTATAAGTTCTGTATGAGACCACAGATCCCCCGCGGGTGCTTTCCAGCT CTTTTT;

### ChrRNA-seq WT and KO

The samples were prepared by isolating chromatin-associated RNA from HEK293T cells in Wild-Type and svtRNA1-2 knock-out conditions in triplicates. The reads were trimmed by Cutadapt (v.1.8.3), discarding processed reads shorter than 10 nt. The processed reads were then mapped to the human reference genome (hg38) using STAR aligner (v.2.5.3a). The basic genome annotation was downloaded from the GENCODE website. The genes were quantified using featureCounts (v.1.6.2) and the counts’ matrix was then normalised using DESeq2. The Gene Ontology enrichment analysis was performed using ShinyGO 0.76[56].

### Miranda target predictions

The sequences of the identified significantly up-regulated genes in Dicer KD, Ago2 KD conditions were downloaded from Biomart[57]. The identified svtRNA1-2 target genes were predicted by running miRanda with the following settings -sc 150 -en -30 -strict (v.3.3a)[31]. The information such as gene target name mapped svtRNA name, its coordinates and the scores were obtained from the miRanda output file. The annotation of the target genes was obtained from Biomart. The custom gene feature file (Intron, Exon, UTR5, UTR3, UTR5/Exon meaning UTR Exon junction same for UTR3/Exon) in BED format was constructed from GENCODE basic annotation file. The canonical transcript file was constructed from the GENCODE basic annotation file taking one transcript per gene with the highest number of relevant tags that represent transcript confidence. Next, we used bedtools (v.2.27.10) intersect function with selected options: -wa, -wb, -loj to identify the features of the mapped vault RNA fragment. By the intersection of predicting mapping coordinates and the genomic features coordinates file. Then we used Python 3 scripts to visualise the result. The calculated percentage of predicted mappings in a particular feature was done by normalising the number of predicted mappings to the feature’s length.

### PhastCons Conservation Analysis

We used phastCons scores resulting from the multiple alignment of 100 vertebrate genomes to the human genome. The files were downloaded in wigFix format from the UCSC database (hg38.100way.phastCons). We extracted the phastCons scores corresponding to vtRNA1-2 and calculated the average score. The density of the normalised phastCons scores was visualised using python3.

### Secondary and tertiary structures of vtRNA molecules

We used RNA immunoprecipitation pull-down selective 2’-hydroxyl acylation followed by Primer Extension (icSHAPE-MaP)[25] dataset to refine vtRNA secondary and tertiary structure predictions. RNAfold[58] command line version with vault RNA sequences in FASTA format and additional -- shape=vault RNA icSHAPE-MaP data were used to predict secondary structures.

>vtRNA1-1

GGGCUGGCUUUAGCUCAGCGGUUACUUCGACAGUUCUUUAAUUGAAACAAGCAACCUGUCUGGGUUGUUCGAGACCCGCGGGCGCUCUCCAGUCCUUUU

(((((((…..((((.((((((…((((…(((…..)))….(((((((….))))))))))))).))))))))…..)))))))…. (−58.45)

>vtRNA1-2

GGGCUGGCUUUAGCUCAGCGGUUACUUCGAGUACAUUGUAACCACCUCUCUGGGUGGUUCGAGACCCGCGGGUGCUUUCCAGCUCUUUU

(((((((….(((((.((((….((((…..…..((((((((….))))))))))))..)))).)).)))..)))))))…. (−53.30)

>vtRNA1-3

GGGCUGGCUUUAGCUCAGCGGUUACUUCGCGUGUCAUCAAACCACCUCUCUGGGUUGUUCGAGACCCGCGGGCGCUCUCCAGCCCUCUU

(((((((…..((((.((((…..…..(((.((.(((.(((……))).))).)))))))))))))…..)))))))….(−58.21)

There were no icSHAPE-MaP data for vault RNA2-1.

Next, we used RNAComposer Automated RNA Structure 3D Modeling Server[59] to model 3D structures of the vtRNA. The output structure was then downloaded in .pdb format and imported into Jmol 9 (http://www.jmol.org/) for visualisation and further modifications.

## Acknowledgements

We thank to all members of the Gullerova group. We acknowledge the ENCODE Consortium and the ENCODE production laboratory(s) for generating the dataset(s). This work was supported by the Senior Research Fellowship by Cancer Research UK [grant number BVR01170] awarded to M. G., EPA Trust Fund [BVR01670] awarded to M.G. and Lee Placito Fund awarded to M.G.

The ERBB4 target explorer was generated using QIAGEN Ingenuity Target Explorer (QIAGEN, Inc., https://targetexplorer.ingenuity.com/).

The experimental illustrations were created with BioRender.com

## Author contributions

A.A., R.F.K., A.D.F. and M.G. performed the experiments and analysed the data. J.T. conducted all bioinformatics analyses under I.C. supervision. M.G. conceived, designed and supervised the study and wrote the final draft of the manuscript.

## Data availability

Information about all published data sets used in this study is summarised in Supplementary Table 4.

Raw data are deposited in GEO accession number GSE198298 and accessible with token: gtczyqgcvlmntkz (https://www.ncbi.nlm.nih.gov/geo/query/acc.cgi?acc=GSE198298).

ChrRNA-seq data from WT and vtRNA1-2 KO cells can be viewed here: https://genome-euro.ucsc.edu/s/Janca9/vtRNA1%2D2_KO_chrRNA%2Dseq

## FIGURE LEGENDS

**Figure S1.**
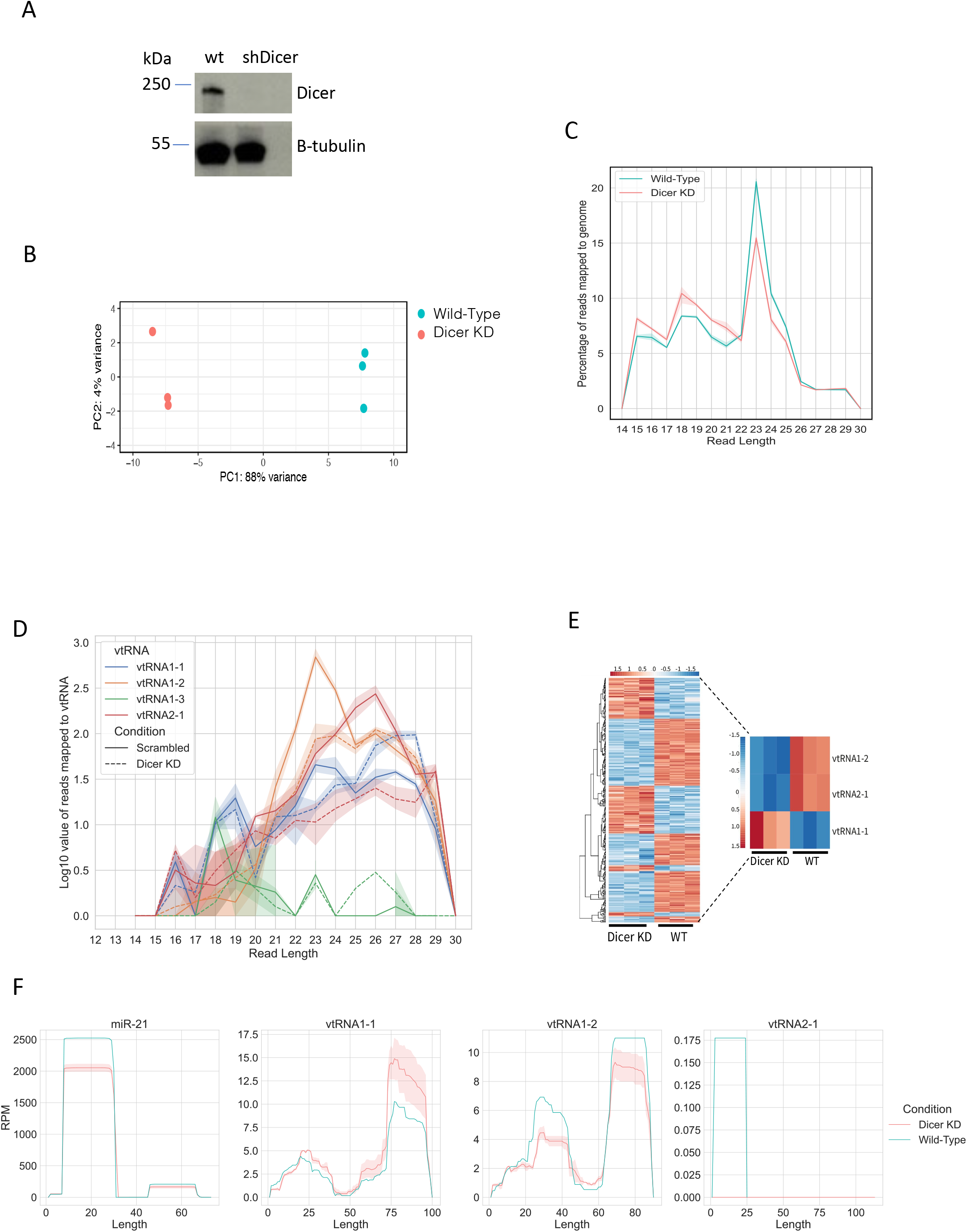
**A)** Representative Western blot showing the Dicer protein levels in WT and Dicer KD cells. Samples were collected 3 days after addition of doxycycline. **B)** Principal-component analysis (PCA) plot shows three biological replicates corresponding to the samples from HEK293T cells in WT and Dicer KD conditions. The RNA composition of cells in the WT condition differed significantly from those in the Dicer KD condition. **C)** Size distribution of mapped reads to the human genome in Dicer KD and WT conditions. **D)** Size distribution of mapped reads to the vault RNA in Dicer KD (dashed line) and WT (full line) conditions. The shaded area around the line represents the standard deviation of the mean. **E)** Heatmap showing small RNA-seq data upon knock-down of Dicer in HEK293T cells. The colour scale indicates normalized intensities (z-score). The heatmap contains all genes that were significantly altered (adjusted p < 0.05) upon Dicer KD. Zoomed heatmap shows significantly expressed vtRNA genes. **F)** Per base read coverage of vtRNA. The x-axis shows the length of transcripts, and the y-axis shows the normalised number of mapped reads in RPM (Reads Per Million). miR-21 acts as a positive control.

**Figure S2.**
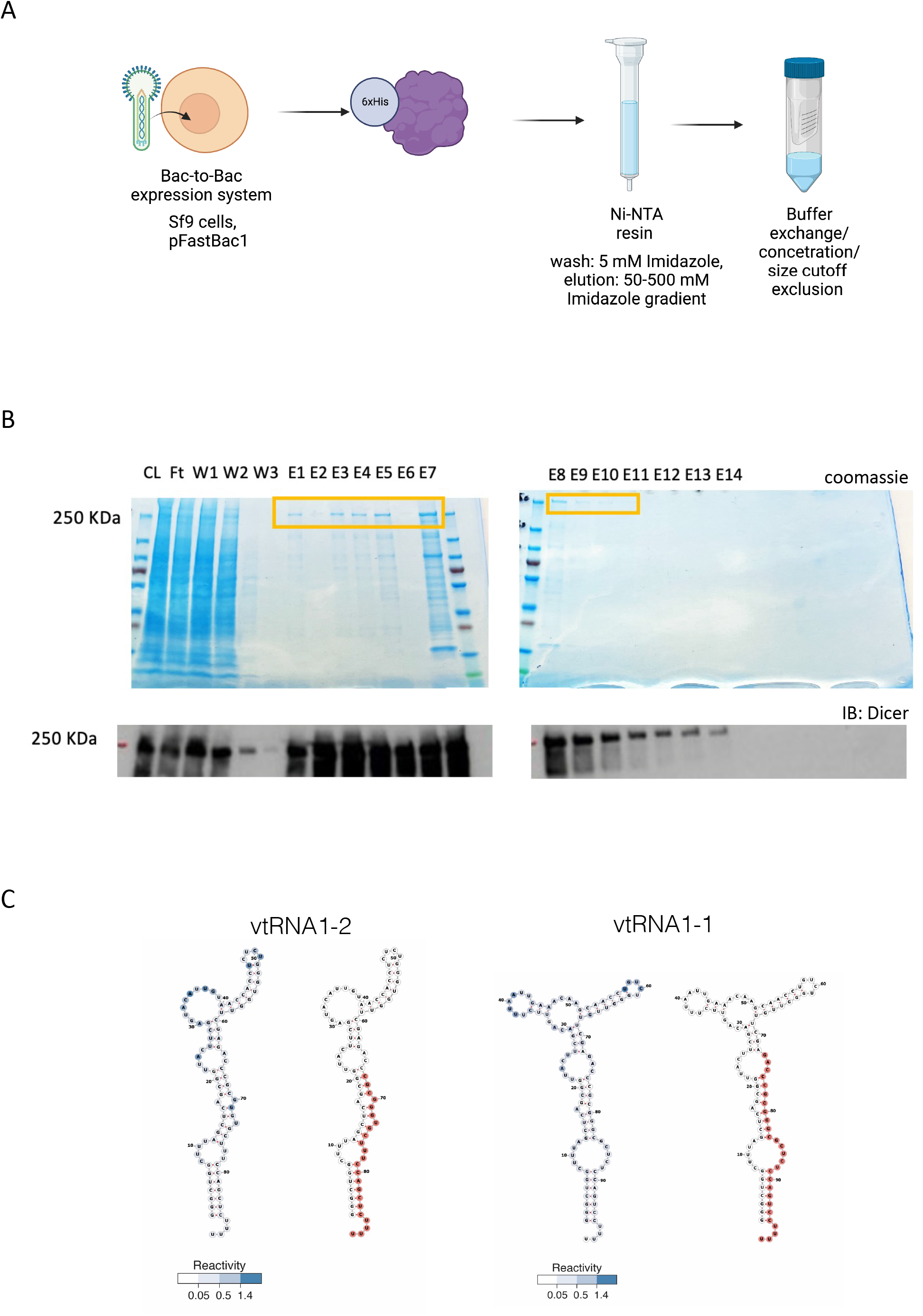
**A)** Diagram summarising 6xHis-Dicer purification strategy from insect cells. **B)** Representative protein gel images showing fractions with purified Dicer protein stain by Coomassie (upper image) or detected by Western blot using anti-Dicer antibody (lower image), (CL=cell lysate, FT= flowthrough, W= washes, E=elution). **C)** Predicted secondary structures of vtRNA1-1 and 1-2. Reactivity score is shown in blue.

**Figure S3.**
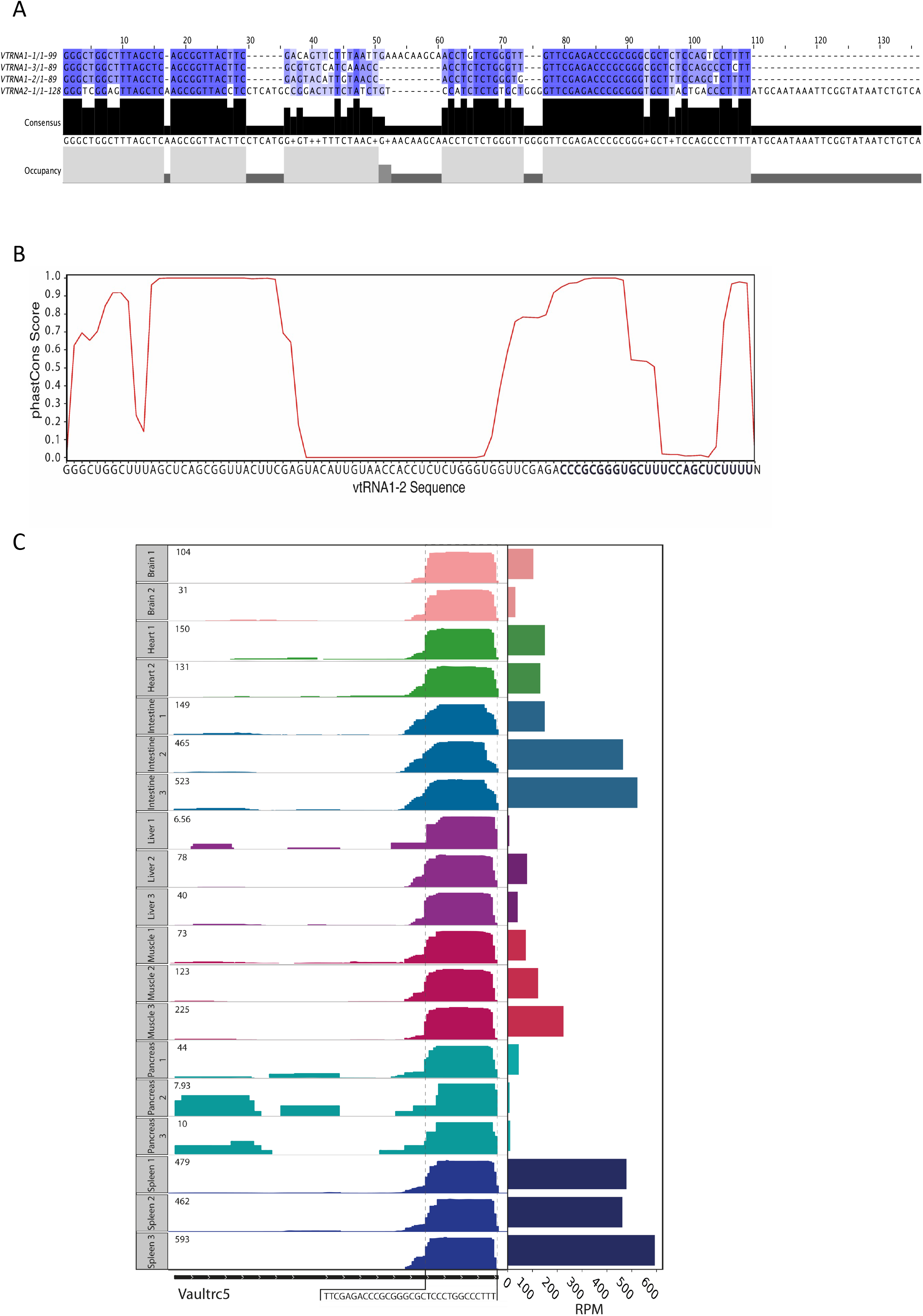
**A)** Vault RNA sequences alignment showing high sequence similarity. **B)** Conservation analysis of vtRNA1-2. The x-axis shows the sequence of vtRNA1-2, and the y-axis shows the conservation scores calculated from the phastCons multiple alignments of 100 vertebrate species. **C)** The IGV tracks showing coverage plots for mouse vault RNA gene (Vaultrc5) from different tissues (Brain, Heart, Intestine, Liver, Muscle, Pancreas, Spleen) in duplicates/triplicates. The tissues are colour-coded and the expression is shown as RPM in the top left corner. The black rectangular at the bottom displays the length of the gene, and the white arrows represent the strand identity. The framed sequence shows the identified 3’ end fragment of Vaultrc5.

**Figure S4.**
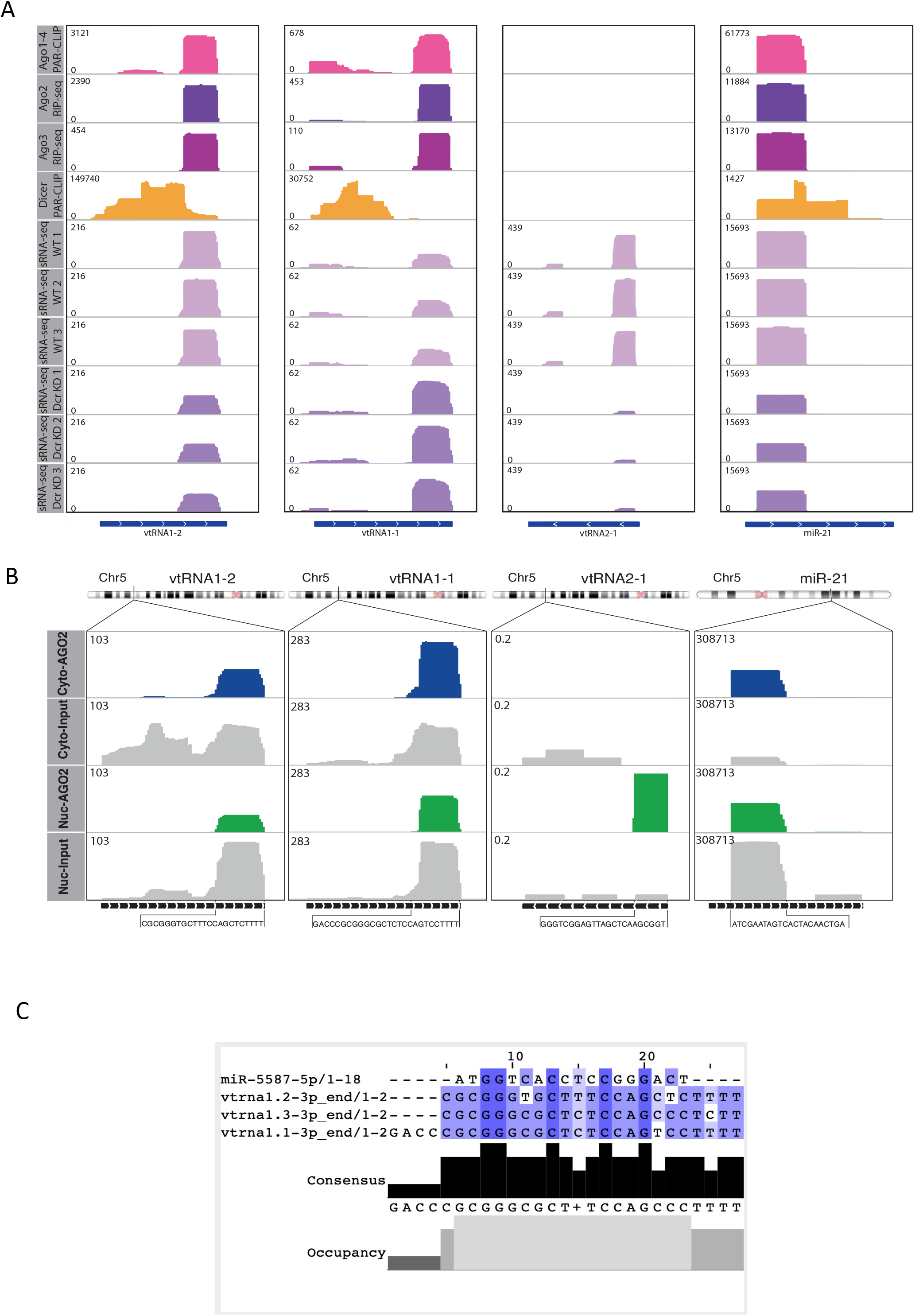
**A)** IGV tracks of AGO1-4 PAR-CLIP, AGO2/3 RIP-Seq, Dicer PAR-CLIP and small RNA-seq in Dicer KD and WT conditions. Data values are shown in the top left corner. miR-21 as a positive control. The bottom line indicates the length of the transcript. **B)** IGV tracks showing reads mapping to vtRNA in nuclear and cytoplasmic AGO2 PAR-CLIP data. The bottom black line indicates the length of the transcript, and the sequence below represents the interacting fragment. The top chromosome ideogram shows the position of each gene. miR-21 acts as a positive control for AGO2 binding. **C)** svtRNA and miR-5587 sequence alignment show bases with high sequence similarity.

**Figure S5.**
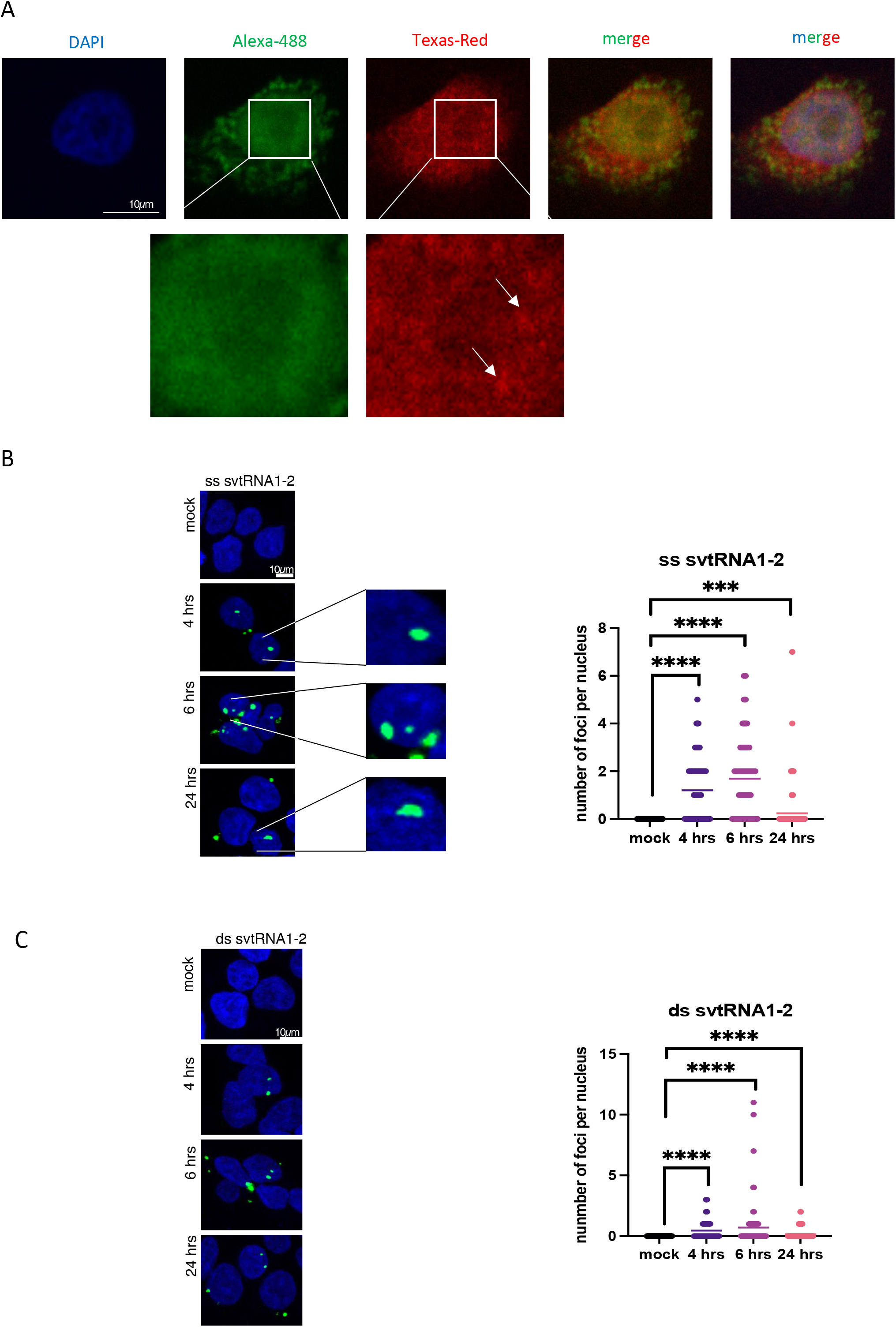
**A)** Representative confocal images showing fluorescence of probes 1 (green) and 2 (red) as in Figure 2, in HeLa cells. Zoomed parts are depicted in white squares, clusters are indicated by arrows. DAPI was used to stain nuclei. **B)** Confocal microscopy images show nuclear localisation of double-stranded (ds) svtRNA1-2 (green) at different time points. DAPI was used to stain nuclei of the cells. Images were quantified using Cell profiler software. Significance was calculated using t-test (****p< 0.0001; ***p< 0.001). **C)** Confocal microscopy images show nuclear localisation of single-stranded (ss) svtRNA1-2 (green) at different time points. DAPI was used to stain the nuclei of the cells. Images were quantified using Cell profiler software. Significance was calculated using t-test (****p< 0.0001; ***p< 0.001).

**Figure S6.**
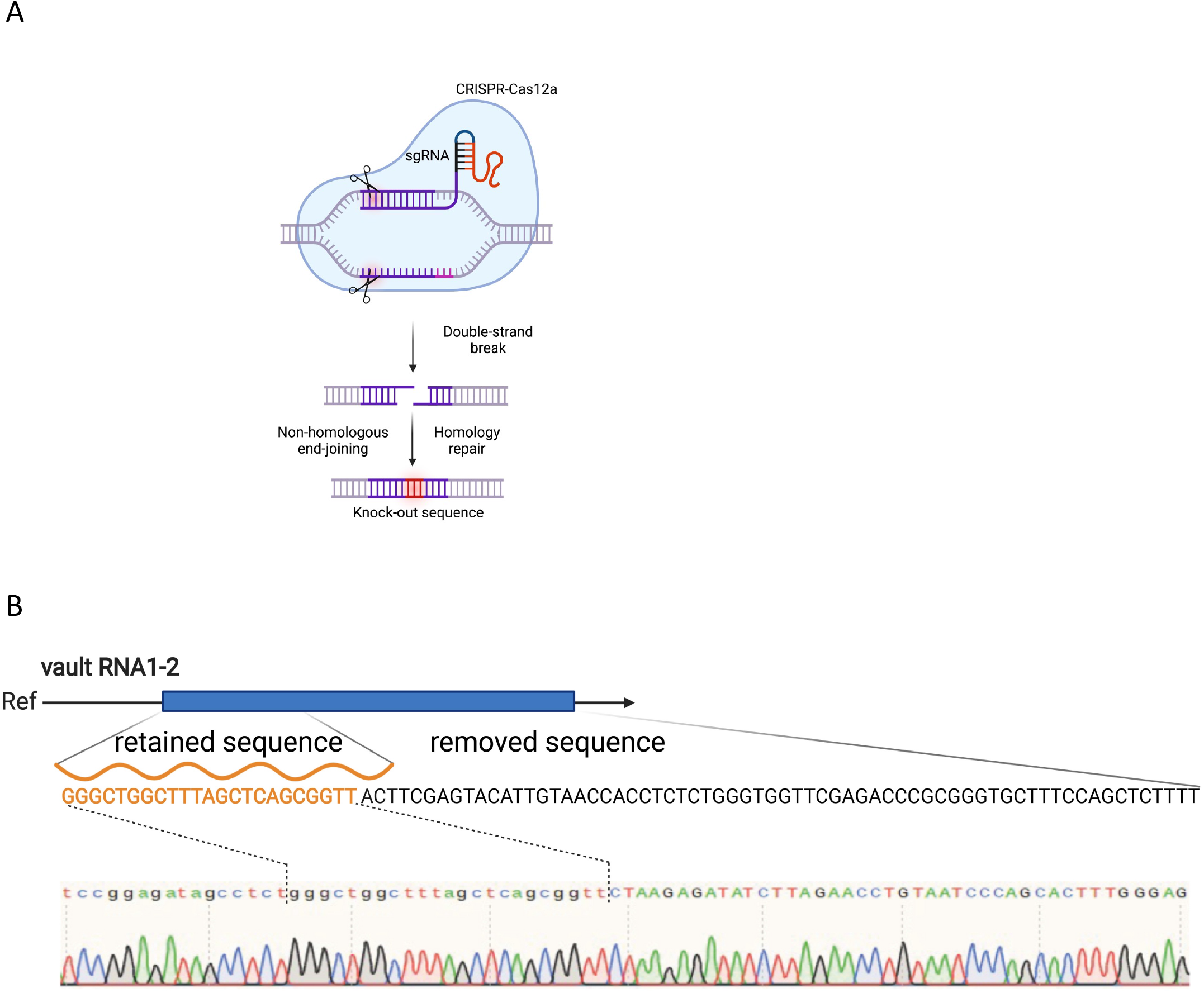
**A)** Schematic representation of CRISPR/Cas12a strategy. **B)** Sanger sequencing results showing sequence of vtRNA1-2 in vtRNA1-2 knock-out. Retained and removed parts of vtRNA1-2 gene are highlighted.

**Figure S7.**
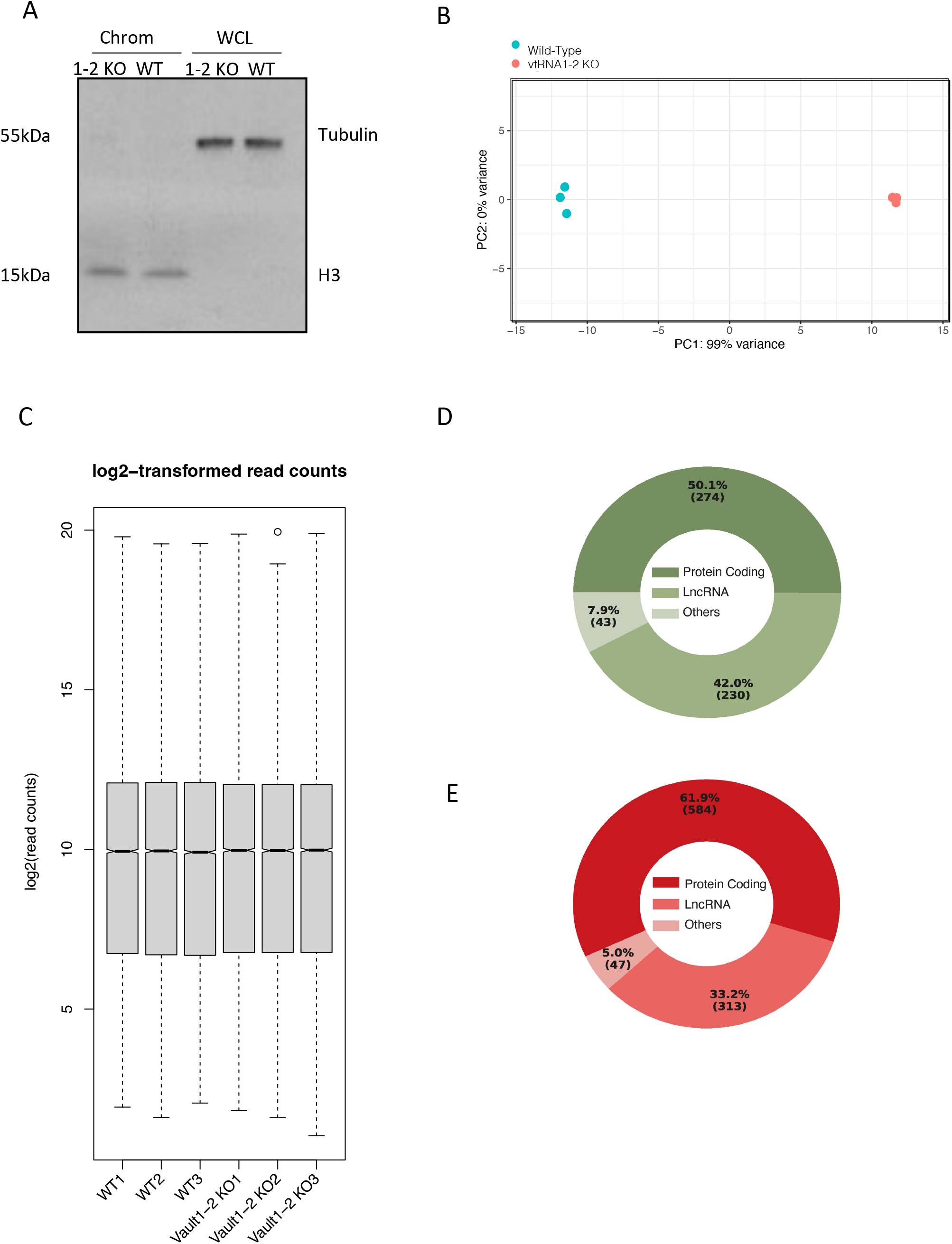
**A)** Western blot of chromatin and whole cell lysate fractions components isolated for ChrRNA sequencing. **B)** Principal-component analysis (PCA) plot shows three biological replicates corresponding to the samples from HEK293T cells in WT and vtRNA1-2 KO conditions. The RNA composition of cells in the WT condition differed significantly from those in the vtRNA1-2 KO condition. **C)** Box plot showing log2-transformed read counts in 3 biological replicates per condition. **D)** Pie chart showing gene type distribution of down-regulated genes in vtRNA1-2 KO cells. **E)** as in **D**, for up-regulated genes

**Figure S8.**
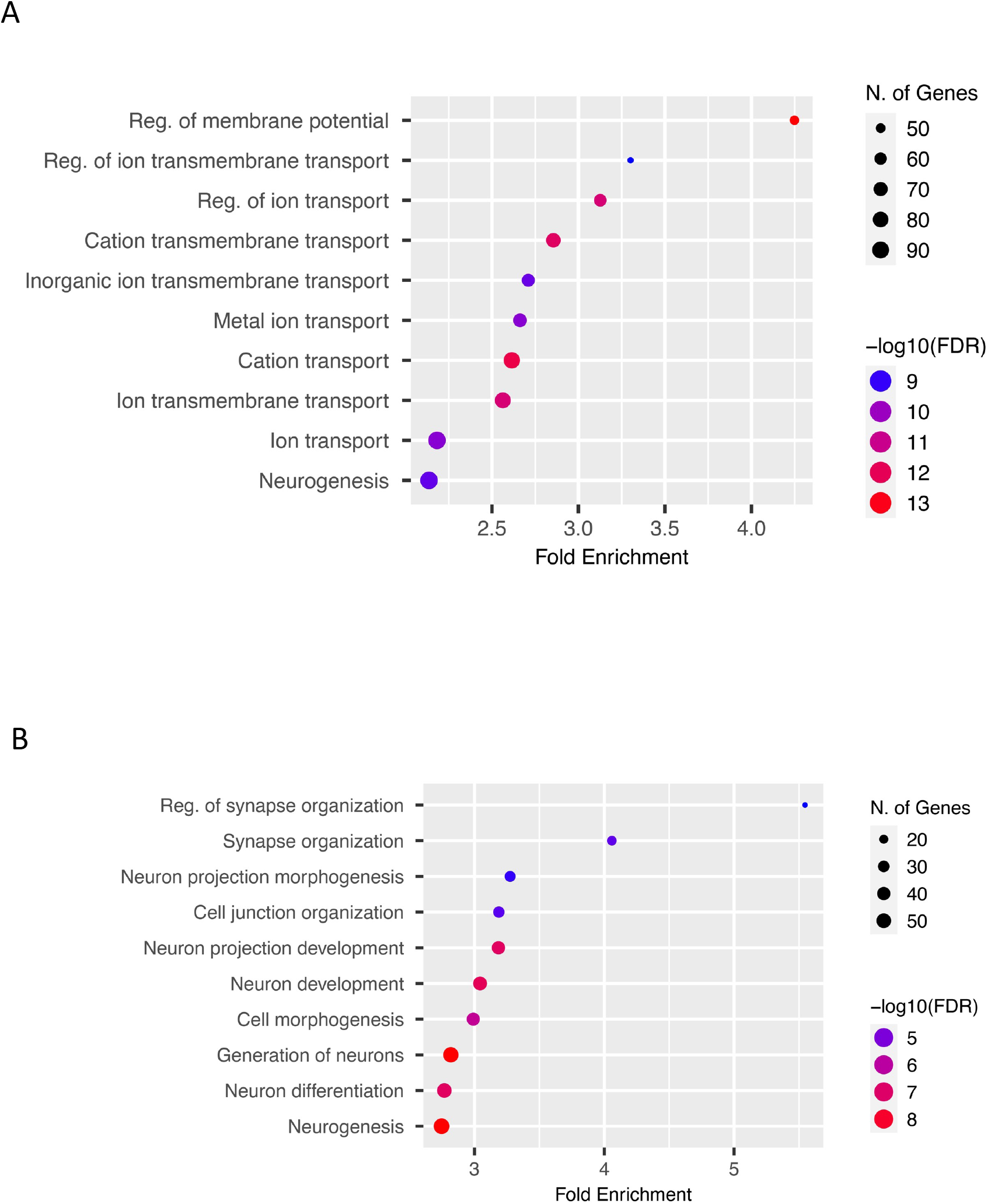
**A)** GO analysis showing up-regulated biological processes in vtRNA1-2-KO when compared to WT cells. Bars represent the −log10 (P-value) and the frequency polygon (black line) denotes the number of genes. **B)** As in **A**, for down-regulated genes.

**Figure S9.**
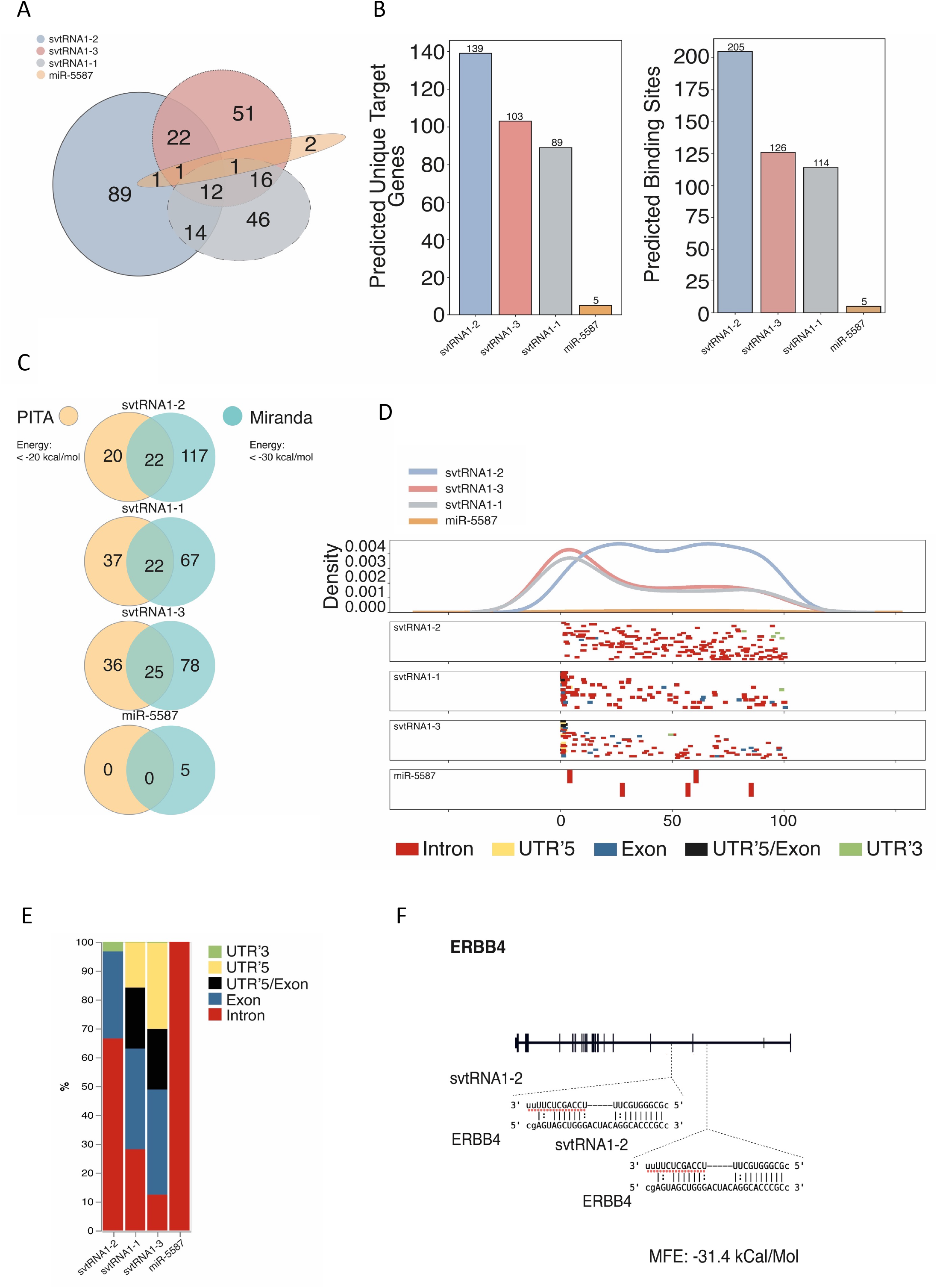
**A)** Venn diagram showing an overlap of target genes predicted for each svtRNA and miR-5587. **B)** Bar chart showing predicted unique target genes (left) and binding sites (right) for each svtRNA and miR-5587. **C)** Pie chart comparing a number of predicted target genes identified by PITA and MiRanda software for each svtRNA and miR-5587. **D)** Distribution of predicted svtRNA binding sites mapped along normalised target transcripts length and their corresponding gene features. **E)** svtRNA binding site counts normalised to the length of a feature displayed in overall percentage. **F)** Schematic representation of ERBB4 gene with predicted binding sites for svtRNA1-2.

**Figure S10.**
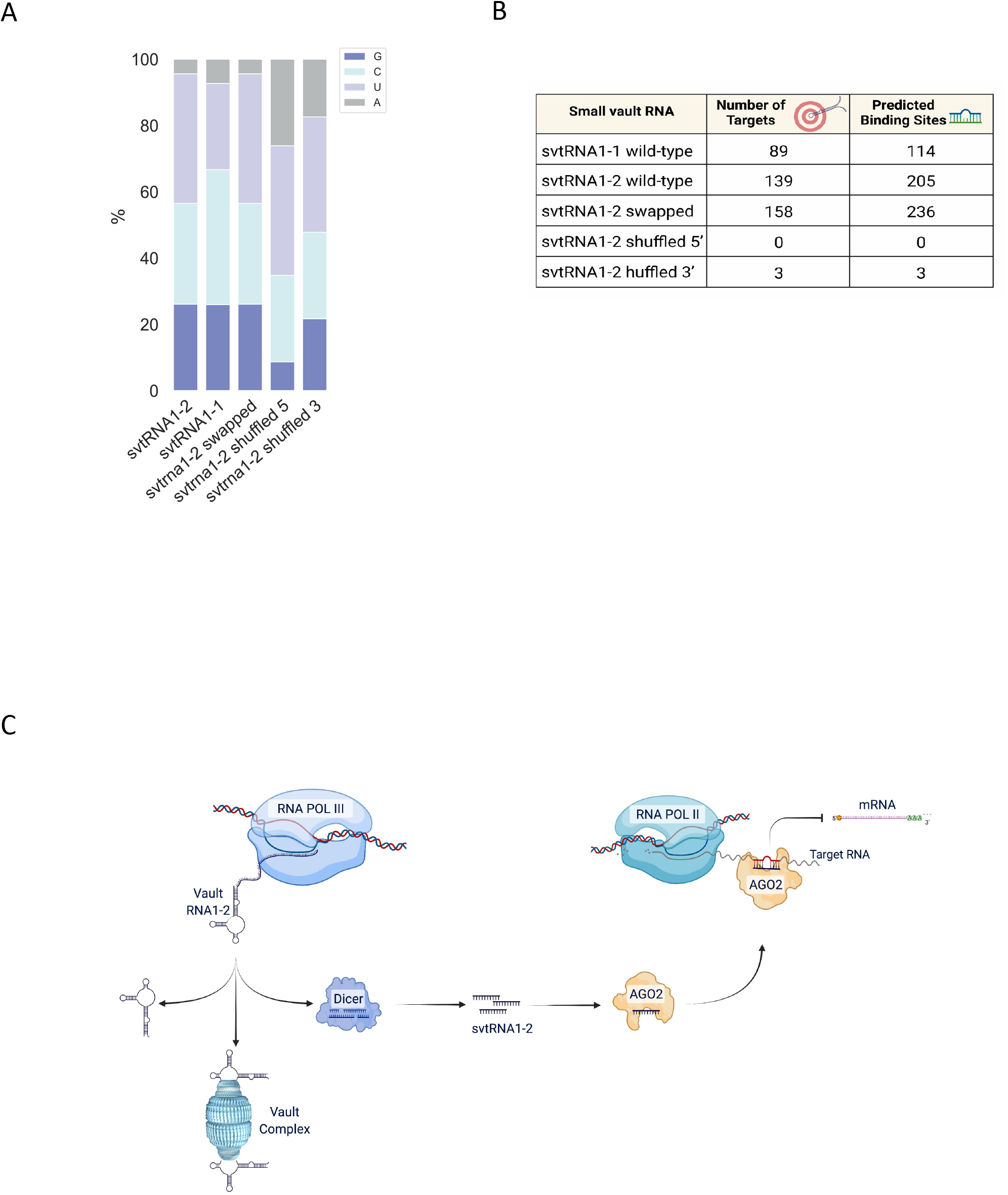
**A)** The bar chart represents the base composition of each sequence displayed as a percentage. **B)** Table reporting number of predicted targets and number of predicted binding sites per sequence. **C)** The model of svtRNA1-2 nascent RNA silencing.

